# Sensory and decisional components of endogenous attention are dissociable

**DOI:** 10.1101/231613

**Authors:** Sanjna Banerjee, Shrey Grover, Suhas Ganesh, Devarajan Sridharan

## Abstract

Endogenous attention acts by enhancing sensory processing (perceptual sensitivity) and prioritizing gating of attended information for decisions (choice bias). It is unknown if the sensitivity and bias components of attention are under the control of common or distinct mechanisms. We tested human observers on a multialternative visuospatial attention task with probabilistic cues, whose predictive validity varied across locations. Analysis of behavior with a multidimensional signal detection model revealed striking dissociations between sensitivity and bias changes induced by cueing. While bias varied in a graded manner, reflecting cue validities, across locations, sensitivity varied in an ‘all-or-none’ fashion, being highest at the cued location. Cue-induced modulations of sensitivity and bias were uncorrelated within and across observers. Moreover, bias changes, rather than sensitivity changes, covaried robustly with key metrics of reaction times and optimal decision-making. Our results demonstrate that endogenous attention engages not a unitary process, but dissociable sensory and decisional processes.

## Introduction

Attention is the remarkable cognitive capacity that enables us to select and process only the most important information in our sensory environment. In laboratory tasks, endogenous attention is engaged by cues that are predictive of upcoming stimuli or events of interest. Numerous studies have explored the behavioral and neural correlates of endogenous cueing of attention, and have reported systematic effects both on sensory processing (Bashinski & Bacharach, 1980; Cohen & Maunsell, 2009; Luck et al, 1996; McAdams & Maunsell, 1999; Ungerleider, 2000) and decision-making (Moran & Desimone, 1985; Baruni et al 2015; Müller & Findlay, 1987; Shaw, 1984). Yet, it remains actively debated whether sensory and decisional effects of endogenous cueing are controlled by a common, unitary attention mechanism or distinct, independent mechanisms (Buschman & Kastner, 2017; Eckstein et al, 2013; Krauzlis et al 2014; Luck et al, 1996; Luo & Maunsell, 2015).

Signal detection theory (SDT), a highly successful framework for the analysis of behavior (Green & Swets, 1966; Ashby, 1992; Macmillan & Creelman, 2005; Carandini & Churchland, 2013; Cohen & Maunsell, 2009; Hawkins et al 1990; Pestilli et al, 2011; Shaw, 1984), specifies two key psychophysical metrics to quantify the sensory and decisional effects of endogenous cueing: i) perceptual sensitivity (d’), which measures the improvement in the quality (SNR) of sensory information processing of the attended stimulus and ii) choice bias (b), which measures the relative weighting of information from the attended stimulus for guiding behavioral decisions (Buschman & Kastner, 2015; Carandini & Churchland, 2013)

A rich literature has reported diverse, and often contradictory, effects of endogenous cueing on sensitivity and bias. For instance, past studies have reported a benefit (Bashinski & Bacharach, 1980)or no benefit (Müller & Findlay, 1987; Shaw, 1984) for sensitivity at the cued location, a cost (Downing, 1988; Hawkins et al, 1990; Luck et al, 1996) or no cost (Bashinski & Bacharach, 1980) for sensitivity at uncued locations, and graded (Müller & Findlay, 1987) versus no modulation of bias (Bashinski & Bacharach, 1980) across locations. Studies investigating the neural correlates of sensitivity and bias have also produced conflicting findings (Luo & Maunsell, 2015, 2018; Baruni et al, 2015).

A key reason for these contradictions is the lack of an appropriate psychophysical (SDT) model for analyzing behavioral responses in endogenous cueing tasks of the type shown in Figure 1A, which are ubiquitous in attention literature (Bashinski & Bacharach, 1980; Cohen & Maunsell, 2009; Corbetta et al, 2000; Lovejoy & Krauzlis, 2009, 2017; Luo & Maunsell, 2015; Müller & Findlay, 1987; Wang & Krauzlis, 2018; Zénon & Krauzlis, 2012). In such tasks, the observer is cued to attend to one of two (or multiple) stimulus locations. At a random time following cue onset an event of interest, for example a change in grating orientation, occurs at one of the locations (“change” trials). On some trials, no events (changes) occur at any location (“catch” trials). The observer must detect and report the location of the change (change trials) or indicate that no change occurred (catch trials). These tasks typically employ probabilistic, spatial cues, whose predictive validity varies across locations, and do not employ post-hoc response probes (Rahnev et al, 2011; Wyart et al, 2012). Unlike response probe tasks, however, such tasks enable measuring a competitive spatial choice bias across locations. On each trial, the subject makes a single choice, by comparing sensory evidence across multiple locations, to detect and localize the change. Subjects’ responses can be analyzed with SDT, in principle, to quantify spatial choice bias to each location.

**Figure 1.**
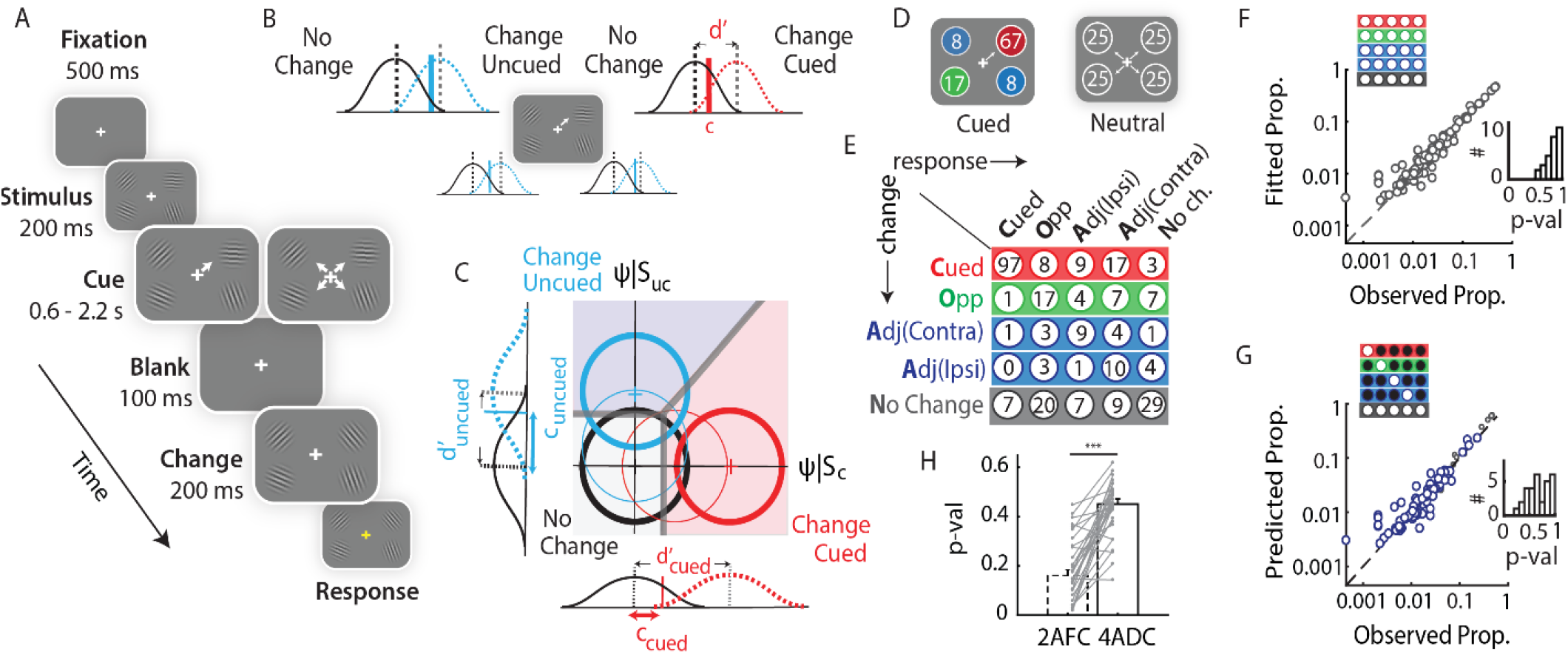
Evaluating a model for analyzing behavior in standard attention tasks. **A**. A “probabilistically cued attention task involving change detection. After fixation (500 ms), four Gabor patches appeared (200ms) followed by an attentional cue (central arrow). After a variable delay (600-2200 ms; exponentially distributed) all patches disappeared briefly (100 ms). Upon reappearance, either one of four patches changed in orientation (“change” trials) or none changed (“no change” trials). After 200 ms the fixation cross changed to yellow, instructing the subject to indicate the location of change, or no change, by pressing one of five different buttons. **B**. Multiple one-dimensional signal detection models representing independent Yes vs. No (change vs. no change) decisions at each location. Black Gaussians: noise distribution for “no change” trials. Red and blue Gaussians: signal distribution for “change” trials at the cued location and uncued location, respectively. Dashed vertical lines: means of signal and noise distributions, whose difference quantifies sensitivity (d’) at each location. Solid vertical lines: Criterion (detection threshold) at each location (c). **C**. Multidimensional (m-ADC) signal detection model, schematized here for changes at the cued and an uncued location. Orthogonal axes: Decision variables at each location. Circles: Gaussian decision variable distribution for “change” trials at the cued location (red); uncued location (light blue); and ‘no change’ trials (black). Thin and thick contours: lower and higher sensitivity values respectively. Thick gray lines: Decision manifold delineating cued, uncued and no change decision domains. Gaussians along the axes: marginal densities at each location. **D**. Proportion of change events at each location in a block with high cue validity (left) and in a block with neutral cues (right). Red: Cued location; green: location opposite to cue (Opp); blue: location adjacent to cue, either in the same visual hemifield as cue (Adj-Ipsi) or in the contralateral one (Adj-Contra). **E**. A contingency table from a representative behavioral session. Rows represent locations of change and columns represent locations of response, both measured relative to the cue. Grey: “No change” trials. Color conventions are as in panel D. (**F-G**) Fitting and predicting responses with a 4-ADC model. F. Response proportions (grey circles) fitted with the 4-ADC model (ordinate) plotted against actual response proportions (abscissa). Data points: response probabilities for each stimulus-response contingency at each angle (data pooled across n=30 subjects). Upper left insets: Subset of stimulus-response contingencies used for fitting. Lower right insets: Distribution of goodness of fit p-values across subjects. **G**. Predicted proportions (blue circles) against actual response proportions by fitting hits and false-alarms alone (see text for details). Small grey circles: fitted proportions. Other conventions are as in panel F. **H**. Goodness-of-fit (randomization test) p-values for observed response probabilities and model predicted response probabilities, for a 2-AFC model (left) and a 4-ADC (right) in a split-half prediction analysis. Grey lines: values for individual subjects.

In practice, however, such tasks present a key challenge for analysis with SDT. Even in their simplest form, such tasks are multi-alternative tasks because they require a choice among three alternatives -- signal event at the cued location, at the uncued location or no signal event -- on each trial. Such tasks have been commonly analyzed with a combination of multiple binary choice (Yes/No) one-dimensional SDT models, with independent decision variables at each location (Fig. 1B) (Bashinski & Bacharach, 1980; Chanes et al, 2013; Luo & Maunsell, 2015; Müller & Findlay, 1987). However, such a formulation is not correct for modeling behavior in multialternative tasks. Because the observer must report a single location of change, modeling behavior with multiple independent Yes/No decisions models produces a choice conflict if a “Yes” decision occurs at more than one location on the same trial (Fig. 1B, see also Discussion).

Here, we analyze human attention behavior in a five-alternative, probabilistically cued attention task with a multidimensional signal detection model, the m-ADC model. This model was recently developed to overcome the pitfalls of analyses with one-dimensional models (Sridharan et al, 2014; 2017) (Fig. 1C). We apply the model, for the first time, to human attention data and test if endogenous cueing engages common or distinct mechanisms for modulating sensitivity and bias. In addition, we develop a novel bias measure that decouples the effects of attention from signal expectation. The results reveal essential dissociations among fundamental components of attention.

## Results

### A model for predicting and fitting behavior in multi-alternative attention tasks

We measured the effect of endogenous cueing of attention on sensitivity and bias by extending the recently developed m-ADC model framework (Sridharan et al., 2014). We developed the model, from first principles, for multi-alternative attention tasks that employ the method of constant stimuli, i.e. in which stimuli can occur at different, unpredictable strengths at each location (Fig. 1C; Figure supplement 1A). This new model is essential, for example, for studies that seek to measure the effect of attention on the psychometric function at cued and uncued locations (Sridharan et al., 2017). The decision manifold in this model comprises a family of intersecting hyperplanes in a multidimensional decision space (Fig. 1C). These hyperplanes are parameterized by decision criteria, which are optimal decision boundaries for distinguishing each class of signal from noise (e.g. change at a given location versus no change), and provide a close approximation to the theoretical optimal decision boundaries for distinguishing signals of one class from another (e.g. changes at one location from another, Figure Supplement 1D-E). The model is able to quantify perceptual sensitivities and decision criteria from stimulus-response contingency tables for tasks with any number of alternatives.

First, we fit a 4-ADC model for each subject’s 5×5 stimulus-response contingency table obtained from the five-alternative task (exemplar table in Fig. 1E). Because changes could occur at one of 6 orientation change values, the contingency table for each subject contained 63 independent observations (details in Methods). For these analyses, responses across two of the uncued locations (ipsilateral and contralateral) were averaged into a single contingency (adjacent) because responses were not significantly different across these locations (p>0.2 for hits, misses and false alarms at these locations, signrank test). This simplification significantly reduced the number of model parameters to be estimated (Methods). Goodness-of-fit p-values obtained from a randomization test (based on the chi-squared statistic) were generally greater than 0.7 (median: 0.84; range: 0.57-0.98; Fig. 1F, lower inset), indicating that the model successfully fit observers’ responses in this multialternative attention task.

Next, we tested the model’s ability to predict individual subjects’ responses: for this we fit the model using only a subset of the observers’ behavioral choices (33%-58%) and tested its ability to predict their remaining (42%-67%) choices (Fig. 1G). Three different subsets of contingencies were selected for fitting: i) hits and false alarms (Fig. 1G), ii) false-alarms and misses (Figure Supplement 1B) or iii) hits and misses (Figure Supplement 1C). Correct rejection responses were included either implicitly (cases i and ii) or explicitly (case iii; see Methods for details). The model was able to predict all of the remaining observations in the contingency table with high accuracy (Fig. 1G, Figure Supplement 1B-C); p-values of randomization fit for predictions (median [95% CI]) case i: 0.69 [0.58-0.87]; case ii: 0.33 [0.25-0.48]; case iii: 0.60 [0.50-0.75]).

Finally, we tested whether a single, unified 4-ADC model provided better fit to the data than a combination of 2-AFC models for modeling decisions at each location (Chanes et al, 2013; Luo & Maunsell, 2018), (Fig. 1H). We performed a split-half analysis by dividing data from each subject’s experimental session into two halves. First, we evaluated how consistently each model estimated sensitivity and bias across the two halves of the experiment, by correlating the parameters estimated from the first half with parameters estimated from the second half. We observed stronger correlations (higher rho values) for the 4-ADC estimates as compared to 2-AFC estimates, in all but one case. Next, we performed a prediction analysis based on the same split-half analysis of the data by correlating the model-predicted response probabilities from the first half against the observed response probabilities in the second half, and vice versa (Supplementary Methods). We observed consistently higher correlations and stronger R^2^ (coefficient of determination) values across subjects, for the 4-ADC model as compared to the 2-AFC model (median ρ: 2-AFC =0.61, 4-ADC =0.64, median R^2^: 2-AFC =0.74, 4-ADC model-data=0.78; p<0.001; Wilcoxon paired signed rank test). To further confirm this, we performed a randomization test for goodness-of-fit, using the same split-half analysis as described above (see Methods). Again, goodness-of-fit p-values were significantly higher for the 4-ADC model as compared to the 2-AFC model, across subjects (median p goodness-of-fit: data-data =0.16, 4-ADC model-data=0.45, p<0.001; Wilcoxon paired signed rank test; Fig. 1H). These data indicate that the 4-ADC model provided superior fits to the data as compared to a combination of 2-AFC models.

Taken together, these results demonstrate that the 4-ADC model accurately fit observers’ behavioral responses in this five-alternative attention task. To our knowledge, this is the first demonstration of a model capable of predicting human choices in a multialternative attention task.

### Endogenous cueing effects on sensitivity, choice criterion and bias

Does endogenous cueing of attention induce changes of sensitivity, changes of bias or both? Changes were twice as likely at the cued location (66.7%), as all of the other locations combined (opposite: 16.7%, adjacent ipsi/contra: 8.3%, Fig. 1D, left). We tested the effect of these different cue validities on modulations of sensitivity and bias, as estimated by the m-ADC model. Bias was quantified based both on the choice criterion (b_CC_) and on the likelihood ratio (b_LR_) measures (Methods). In the following description, we note the distinction between the terms “criterion” and “choice criterion” (or decision criterion). “Criterion” refers to the “detection threshold”, whose value is measured relative to mean of the noise distribution (e.g. Fig. 1C). The choice criterion, on the other hand, is a measure of bias, whose value represents the deviation of the criterion (detection threshold) from its optimal value (see Methods). Cue validity was not different between cue-ipsilateral and cue-contralateral locations, and parameter values were strongly correlated and not significantly different between these locations (Figure Supplement 2A); these locations were treated as a single cue-”adjacent” location for these analyses (Methods).

First, we tested whether subjects were utilizing information provided by cue regarding the location of the imminent change. We quantified the effect of cueing “raw” performance metrics, including hit, false alarm and correct rejection rates (Fig. 2A). Subjects (n=30) exhibited highest hit rates for changes at the cued location as compared to the other locations, across the range of angles tested (Fig. 2A, top; p<0.001 for difference in hit rates between cued vs. opposite and cued vs. adjacent; Wilcoxon signed rank test, Benjamini-Hochberg correction for multiple comparisons), indicating that subjects were indeed heavily relying on the cue to perform the task. Nevertheless, false-alarm rates were also highest at the cued location (Fig. 2A, bottom right; cued vs. opposite: p=0.0014; cued vs. adjacent: p<0.001). We asked whether this pattern of higher hit rates concomitantly with higher false alarm rates occurred due to a higher sensitivity at the cued location, a higher bias (lower choice criterion) at the cued location, or both (Macmillan & Creelman, 2005; Steinmetz & Moore, 2014), (Methods), by estimating these psychophysical parameters with a 4-ADC model.

**Figure 2.**
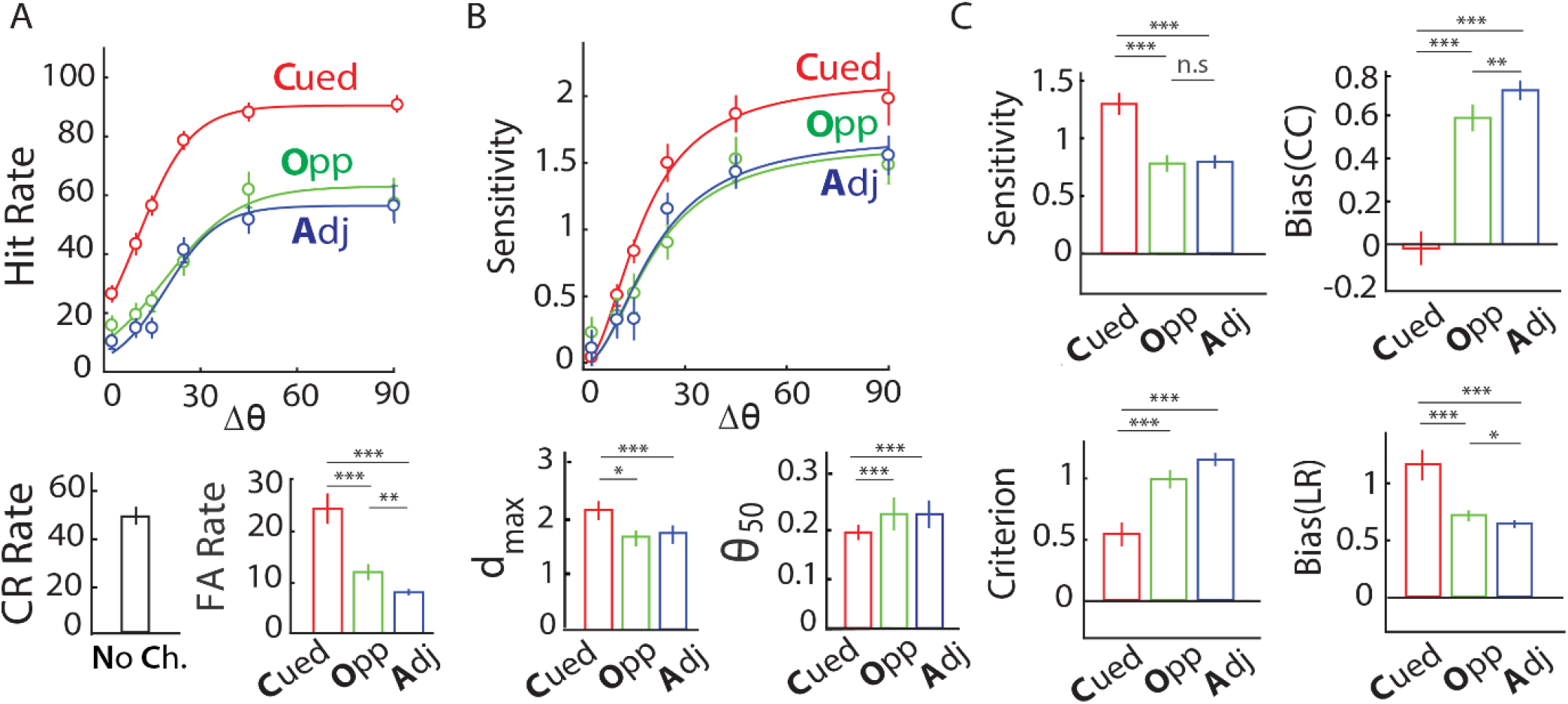
Sensitivity and bias changes induced by endogenous cueing in a multialternative attention task. **A**. (Top) Average psychometric function (n=30 subjects) showing hit rates across different change-angles (Δθ). Curves: Sigmoid fit. (Bottom) Mean correct rejection (left) and mean false-alarm rates (right). Error bars: jackknife s.e.m. Color conventions are as in Fig. 1B. **B**. (Top) Average psychophysical functions showing sensitivity at different change angles. Curves: Naka-Rushton fit. (Bottom) Asymptotic sensitivity (d_max_, left) and change-angle at half-max sensitivity (Δθ_50_, right). Other conventions are as in panel A. **C**. Median sensitivity (top-left), detection criterion (bottom-left), choice criterion bias (bCC, top-right) and likelihood ratio bias (bLR, bottom-right) at different locations. Error bars: s.e.m. Color conventions are as panel A.

Sensitivity was consistently highest at the cued location as compared to the other, uncued locations (p<0.001; Fig. 2B, top). We fit the 4-ADC model incorporating a parametric form of the psychophysical function (the Naka-Rushton function). Asymptotic sensitivity (d_max_) was highest for the cued location as compared to each of the uncued locations (p<0.001, bootstrap test against a null distribution of differences generated by randomly shuffling location labels; Methods; Fig. 2B, bottom left) whereas change magnitude corresponding to half-max sensitivity (Δθ_50_) was significantly lower for the cued, as compared to each of the uncued locations (p<0.001, bootstrap test; Fig. 2B, bottom right). Similarly, bias was uniformly highest at the cued location compared to the uncued locations: choice criteria were lowest, and likelihood ratio bias highest towards the cued location compared to the other locations (Fig. 2C; p<0.001; corrected for multiple comparisons).

Next, we asked if these modulations of sensitivity and bias reflected a benefit (relative to baseline) at the cued location, a cost at the uncued locations, or both. For this, a subset of the subjects (n=10) who were tested on the cued detection task were also tested on a neutrally-cued version of the task presented in interleaved blocks (Methods). In this task, changes were equally likely (25%) at each of the four locations. Within this pool of subjects, sensitivity for the neutrally cued locations was significantly lower than that at the cued location (p=0.004), but only marginally significantly different from that at the uncued locations (p=0.049; Figure Supplement 2D, top left). On the other hand, biases (b_CC_ and b_LR_) for the neutrally cued locations were intermediate between that of the cued and uncued locations, and robustly significantly different from cued (p=0.002) as well as uncued (p<0.01) locations (Figure Supplement 2D, right).

In sum, these results show that endogenous cueing of attention produced both a higher sensitivity as well as higher bias towards the cued location relative to uncued locations. The increase in sensitivity manifested primarily as a benefit at the cued location. In contrast, cueing produced both a strong benefit in bias at the cued location and a strong cost at uncued locations, relative to baseline.

### Sensitivity and bias: Same or different mechanisms?

Are the enhancements of sensitivity and bias by endogenous cueing mediated by the same or different mechanisms? To answer this question, we evaluated two competing models (Fig. 3A). According to the “common” mechanism model, cue-induced selection biases compete for sensory processing resources to enhance sensitivity, so that sensitivity at each location co-varies systematically with bias at that location. On the other hand, according to the “disjoint” mechanisms model bias and sensitivity modulation are decoupled and independent of each other. We sought to distinguish between these two models by examining several lines of evidence.

**Figure 3.**
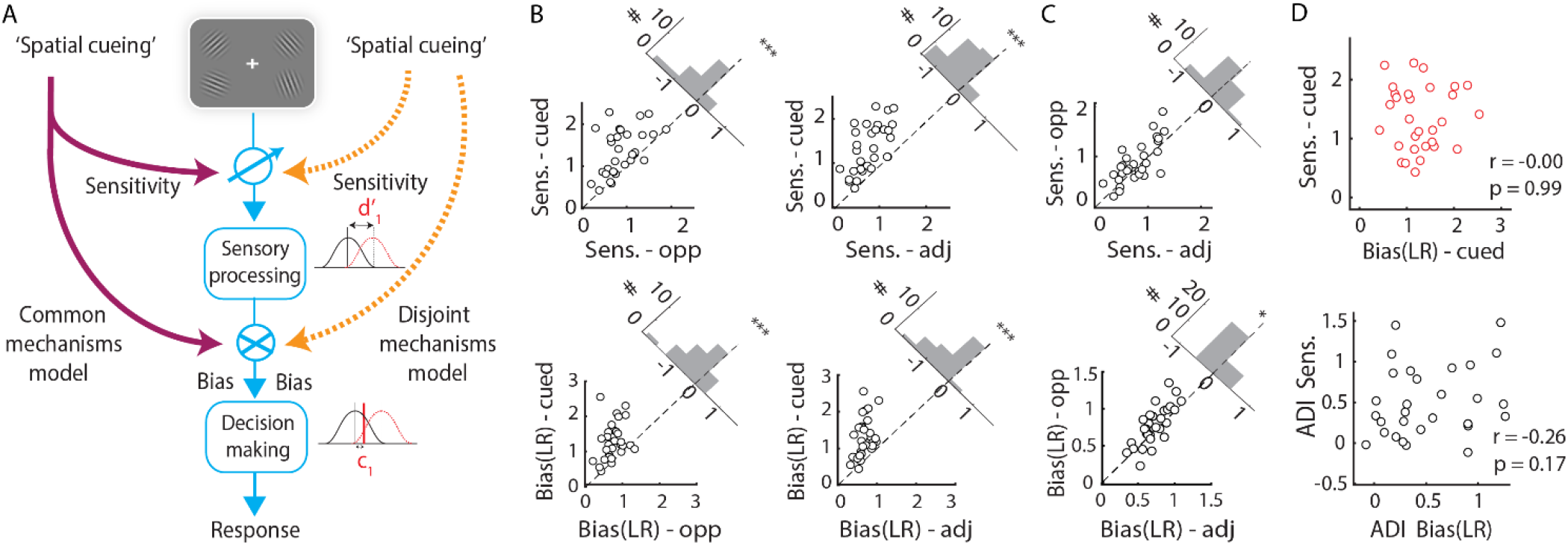
Common versus distinct mechanisms of sensitivity and bias changes. **A.** Schematic of common (left) versus disjoint (right) models by which spatial cueing may modulate sensory processing (sensitivity, d’) and decision-making (bias or criterion). Cueing may modulate sensitivity and bias through a common mechanism (purple arrows) or through distinct mechanisms (orange arrows). **B.** Sensitivity (top) or bias (bottom) at the cued versus opposite (left) or adjacent (right) locations, measured with the 4-ADC model. Data points: individual subjects (n=30). Dashed diagonal line: line of equality. Upper right insets: Histograms indicating difference in parameter values. ***-p<0.001, *-p<0.05 for a difference of medians. **C**. Same as in panel B, but for the opposite versus adjacent locations. Other conventions are as in panel B. **D**. (Top) Covariation between sensitivity and bias at the cued location. (Bottom) Covariation between attention difference indices of sensitivity and bias. Data points: individual subjects.

First, cue validity at the uncued locations occurred at one of two different values (Fig. 1D, left). Of the one third of change trials occurring at the uncued locations, changes were twice as likely at the cue-opposite location (16.7%) as at either of the adjacent locations (8.3% each). We asked if these different cue validities would produce systematically differing values of sensitivity and bias at the different uncued locations. Across the population of subjects (n=30), sensitivities at the opposite location were not significantly different from those at the adjacent locations (p=0.08; Fig. 2C, top left, Fig. 3B-C upper panels). In contrast, choice criterion bias (bCC) was significantly lower for the opposite compared to the adjacent locations (p=0.009; Fig. 2C, top right), and likelihood ratio bias (b_LR_) was significantly higher for the opposite compared to the adjacent locations (p=0.018; corrected for multiple comparisons). Taken together with the graded variation in bias in the neutral cued condition (Fig. 2C, bottom right; Fig. 3B-C, lower panels; Figure Supplement 2D, right), these results indicate that bias, rather than sensitivity, systematically modulated with endogenous cue validity.

We further tested this by evaluating the performance of two modified mADC models, one with sensitivity constrained to be equal across opposite and adjacent locations (m-ADC_eq-d_) and one with criteria similarly constrained (m-ADC_eq-c_). Model selection analysis based on the Akaike Information Criterion (AIC) revealed that the m-ADC_eq-d_ model significantly outperformed both the m-ADC_eq-c_ model and the standard m-ADC model (AIC: m-ADC_eq-d_ = 500.1 [466.2-602.9]; m-ADC_eq-c_ = 508.6 [475.5-614.2]; m-ADC = 508.7 [474.4-614.3]; p<0.001, Wilcoxon signed rank test), whereas there was no significant difference in the performance of the m-ADC_eq-c_ and the standard m-ADC models (p=0.12). The significantly higher evidence in favor of a model that incorporated identical sensitivities, but distinct criteria, at all uncued locations confirms that criteria, rather than sensitivities, varied in a graded manner with endogenous cue validity.

Despite these trends in parameters at the uncued locations, we asked if subjects who exhibited higher sensitivity at the cued location would also exhibit a higher bias at the cued location. Sensitivity and criteria (detection thresholds) were strongly positively correlated at the cued location (ρ=0.6, p<0.001, Figure Supplement 3A) indicating that across observers’ criteria co-varied with sensitivities, in line with a key prediction of the m-ADC model for optimal decisions in this task (Supplementary Methods: Model Derivation, equation 11). On the other hand, neither choice criterion nor likelihood ratio bias were correlated with sensitivity (ρ_cc-d’_ = 0.13 p_cc-d’_ = 0.49;ρ_LR-d’_ = 0.01 p_LR-d’_ = 0.99; Fig. 3D, top and Figure Supplement 3A, last row). The uncued locations (opposite, adjacent) revealed a similar trend of positive correlation between sensitivity and criteria and no correlation (b_CC_) or even a negative correlation (b_LR_) between sensitivity and bias (Figure Supplement 3A). These results demonstrate that although endogenous cueing produced both the highest sensitivity and bias at the cued location, there was no evidence of a positive correlation between these quantities.

Although sensitivity and bias at the cued location did not covary, we asked if subjects who showed the greatest differential sensitivity at the cued location (relative to uncued locations), also showed the greatest differential bias toward the cued location. To answer this question, we tested whether sensitivity and bias modulations, measured either as a difference index (ADI) or a modulation index (AMI) across cued and uncued locations (Methods), were correlated. Neither bias value (choice criterion or likelihood ratio) was significantly co-modulated with sensitivity, as measured with the difference index (ρ_cc-d’_=-0.10, p_cc_ = 0.61; ρ_LR-d’_=-0.26, p_lr_ = 0.17; Fig. 3D (bottom) and Figure Supplement 3B), or with the modulation index (ρ_cc-?’_=-0.25, p=0.19; ρ_LR-d’_=0.25, p=0.18; Figure Supplement 3C). Next, we tested whether sensitivity and bias were co-modulated across experimental blocks in individual subjects. We divided data from each subject’s experimental session into two subsets of contiguous experimental blocks. Difference indices (ADI) or modulation indices (AMI) for bias from each block were subjected to an n-way analysis of variance with the corresponding sensitivity (ADI or AMI) index as a continuous predictor, and subjects as random effects. This analysis revealed that, neither measure of bias was significantly co-modulated with sensitivity, as measured by either modulation index (ADI or AMI), across blocks (p>0.1, for all tests).

Finally, we examined the possibility that the graded variation of bias with cue validity was a consequence of parametric assumptions in the m-ADC model with a “similarity choice model” (Luce, 2005). In this model, sensitivities and biases are estimated, not from a parametric, latent variable model, but by a factoring of the underlying probability densities. Sensitivity and bias parameters estimated by the similarity choice model were strongly correlated with the corresponding m-ADC model parameters (ρ=0.7-0.9, p<0.001), and also exhibited identical trends, including a graded variation in bias with cue-validity and the lack of correlation between sensitivity and bias or their modulations.

To summarize, sensitivity enhancements were strongest at the cued location and not significantly different across uncued locations: bias modulations were more graded across locations and varied systematically with endogenous cue validity. Moreover, modulations of sensitivity and bias by endogenous cueing were uncorrelated across subjects. These results suggest that the “spotlight” model of attention applies primarily to modulations of sensitivity, rather than bias, and overwhelmingly favor the hypothesis that sensitivity and bias changes are mediated by dissociable neural mechanisms in this endogenous attention task (Fig. 3A, right).

### Covariation of sensitivity and bias with reaction times and decision optimality indices

Attention produces systematic effects on perceptual decisions. These effects are typically quantified with reaction times (Eason et al, 1969) and metrics of optimal decision-making (Eckstein et al, 2013). To further disambiguate the “common” from the “disjoint” mechanisms models, we tested if subjects’ reaction times and decision optimality metrics covaried with sensitivity and bias in similar or distinct patterns. Results are reported here for likelihood ratio measure of bias (b_LR_); similar trends were observed for choice criteria (b_cc_).

Reaction times (RT, normalized to cued location; Methods) for all change responses, including both correct and incorrect responses varied in a graded fashion with cue validity; the fastest RTs occurred for changes at the cued, followed by the opposite and adjacent, locations, in that order (p<0.05, corrected for multiple comparisons; Fig. 4A). Similar graded trends were observed when the data were analyzed separately based on hit and false alarm responses (Fig. 4B-C). This graded variation with cue validity suggested a close relationship between RT and bias.

**Figure 4.**
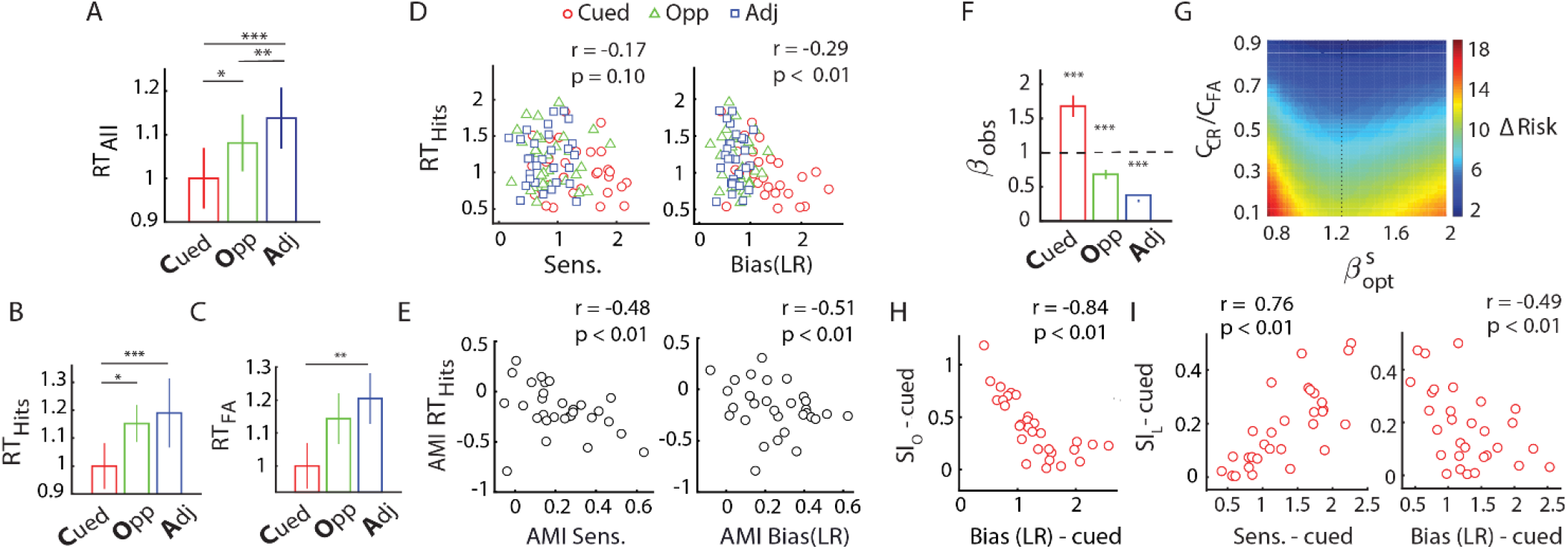
Covariation of sensitivity and bias with reaction times and decision optimality indices. **A.** Normalized median reaction times across subjects for each location (all trials). Color conventions are as in Fig. 1B. **B.** Same as in top panel, but for hit trials. **C.** Same as in top panel, but for false-alarm trials. **D.** Covariation of normalized median reaction times on correct trials for each subject at each location with sensitivity (left) or bias (right). Data points: values for individual subjects at particular locations. Color conventions are as in Fig. 1B. **E.** Same as in panel D, but covariation of the attention modulation indices for median reaction times with those for sensitivity (left) or bias (right). Data points: individual subjects. Other conventions are as in panel D. **F.** Median observed cost ratio (βobs) at the different locations. Dashed line: βopt value (=1) for maximizing correct and minimizing incorrect responses. Color conventions are as in Fig. 1B. **G.** Difference between the actual risk and the optimal Bayes risk (ΔR) for different values of β (x-axis) and different values of the cost ratio of false alarms to correct rejections (C_CR_/C_FA_; y-axis); data pooled across subjects. Warmer colors: larger values of ΔR. Black dots: β* corresponding to minimal ΔR for each C_CR_/C_FA_. **H.** Covariation of the objective sub-optimality index (SI_O_) with bias at the cued location (see text for details). Data points: Individual subjects. **I.** Covariation of the locational sub-optimality index (SI_L_) with sensitivity (left) and bias (right) at the cued location. Other conventions are as in panel E.

Next, we quantified the relationship between RT, sensitivity and bias, considering only correct responses (hits). RT (normalized) and bias were robustly negatively correlated (Fig. 4D, right); ρ=−0.29 p=0.006), whereas RT and d’ were not. (Fig. 4D, left); ρ=−0.17, p=0.10). To test if the RT and bias correlation was rendered more robust by the latent covariation between RT and d’, we performed a partial correlation analysis of RT versus bias controlling for the effect of d’ (and vice versa). We found a significant negative partial correlation between RT and bias, (ρ_p_=−0.21, p=0.04) whereas the partial correlation between RT and d’, controlling for bias, was not significant (ρ_p_=−0.19, p=0.07). We then quantified the co-modulation of RT with d’ and bias across cued and uncued locations. RT AMI was significantly negatively correlated with both bias AMI (ρ=−0.51, p=0.004; Fig 4E, right), and d’ AMI (ρ=−0.48, p=0.007; Fig. 4E, left), but multilinear regression analysis of RT (change angle, average d’ and bias as predictors; Methods) revealed a significantly more negative standardized regression coefficient for RT variation with bias as compared to d’ (β_b-LR_ = −0.082 [−0.21 0.05], β_d’_ = −0.019 [−0.069 0.071] – median [95% CI]; p=0.046, bootstrap test for difference of magnitudes).

We also tested whether the pattern of RT correlations reflected a motoric response bias rather than a cue-induced choice bias. We correlated false alarm rates in each of the six experimental blocks with the mean reaction time on false alarm trials in that block, separately, for the cued and uncued locations. A significant correlation would indicate that faster responses (due to a motor bias) produced more false alarms and, correspondingly, a higher bias. Contrary to this hypothesis, we found no significant correlation between RT and false alarm rates across blocks at any location. (cued: ρ=0.05, p=0.58; opp: ρ=0.13, p=0.18; adj: ρ=−0.15, p=0.10) These results were confirmed with an ANOVA analysis with RT as the response variable and false alarms as continuous predictors with subjects as random factors.

Taken together, these results support a more robust co-variation of reaction times with bias changes, rather than with sensitivity changes, induced by spatial cueing.

Next we tested covariation of sensitivity and bias with metrics of decision optimality. For this, we measured subjects’ observed cost ratio (β_obs_), defined as the ratio of the prior odds ratio to the bias at each location (Supplementary Methods: Model Derivation, equation 12), and compared it with the optimal cost ratio (see Supplementary Methods for details). In the m-ADC model decision rule framework (minimizing risk), β_opt_ equals the ratio of the cost of correct rejections versus false alarms to the cost of hits versus misses (β_opt_ = (C_CR_ − C_FA_)/ (C_Hit_ − C_Miss_); Supplementary Methods: Model Derivation, equation 8). Therefore, subjects with a goal of maximizing successes and minimizing errors in our task should have assumed an optimal cost ratio of unity (β_opt_=1) at all locations (Fig. 4F, dashed horizontal line). Yet, a vast majority of subjects exhibited systematic deviations from this optimum (Fig. 4F): the observed cost ratio was significantly greater than 1 at the cued location, and significantly less than 1 at uncued locations (p<0.001, signed rank test; Fig. 4F), implying that subjects adopted a lower bias than optimal at the cued location and a higher bias than optimal at uncued locations (cost ratio inversely related to bias, Supplementary Methods: Model Derivation, equation 12).

We investigated the reason for this systematic pattern of sub-optimalities. A first, potential scenario is that subjects’ observed cost ratio deviated from the optimal ratio because they perceived a prior signal probability at each location that deviated systematically from the actual signal priors. This scenario arises when subjects fail to detect some proportion of changes, especially when the change in orientation was small. However, this explanation is not tenable because small orientation changes would be difficult to detect at both cued and uncued locations. Hence, the perceived prior ratio would have had to be lower than the actual ratio at all locations. While this scenario can account for the lower than optimal bias at the cued location (Fig. 4F, red bar), it cannot account for the higher than optimal bias at each of the uncued locations (Fig. 4F, green and blue bars). An alternative scenario is that subjects assumed different cost ratios (β-s) at the different locations, such that they judged errors arising from false alarms as more costly compared to errors arising from misses at the cued location (β>1), and vice versa at uncued locations (β<1). Nevertheless, no rational explanation can be readily conceived of for subjects attributing different costs to false alarms and misses at the different locations (Supplementary Methods). Having ruled out these alternative potential scenarios, we propose the following hypothesis: Each subject sought to minimize risk by adopting a “subjective” cost ratio (β^s^_opt_) that was uniform across all locations, but differed from 1 (β^s^_opt_ ≠ 1) and was subjectively optimal to each observer’s own estimate of the relative cost of false alarms and misses. β^s^_opt_ for each observer was measured as the cost ratio (C_FA_/C_CR_; Supplementary Methods) that minimized the difference between the actual risk and the optimal Bayes risk (ΔR^obs-opt^). With data pooled across subjects, β^s^_opt_ occurred at a value of 1.3, across the range of C_FA_/C_CR_ values tested (Fig. 4G)

First, we examined two metrics, which measured the deviation from optimality of each observer’s overall performance: i) an objective sub-optimality index (SI_O_) which measures the deviation of β^s^_opt_ from β_opt_ (Supplementary Methods), and ii) a global sub-optimality index (SI_G_) which measures the deviation of the observed risk from the optimal Bayes risk (ΔR^obs-opt^). SI_O_ was negatively correlated with bias at the cued location (ρ=−0.84, p<0.001; Fig. 4H) but not at the uncued locations (p>0.07), whereas it was not correlated with sensitivity at any location (p>0.4). Complimentarily, SI_G_ was positively correlated with bias and negatively correlated with sensitivity at both opposite and adjacent locations (SI_G_ vs. bias: ρ_opp_=0.71, p_opp_<0.001; ρ_adj_=0.81, p_opp_<0.001, Figure Supplement 4B. SI_G_ vs. sensitivity: ρ_opp_=−0.61, p_opp_<0.001; ρ_opp_=−0.39, p_opp_=0.03) but not at the cued location (p=0.153). Next, we quantified deviation from optimality at each location by defining a locational sub-optimality index (SI_L_) which measures the deviation of the cost ratio at each location (βobs) from the individual’s optimal cost ratio (β^s^_opt_)). SI_L_ was significantly lower (subjects more optimal) at the cued location than at either uncued location (p<0.001; Figure Supplement 4A). SI_L_ was negatively correlated, across subjects, with bias at the cued location, (ρ=−0.5, p=0.006), whereas it was positively correlated with sensitivity (ρ=0.76, p<0.001; Fig. 4I).

To summarize, subjects who exhibited a greater bias toward the cued location, and lower bias toward uncued locations, made more optimal decisions overall in this attention task (Fig. 4H-I; Figure Supplement 4A). Decisional optimality was highest at the cued location and showed opposite patterns of correlations with sensitivity and bias (Fig. 4H-I).

Taken together, this dissociation between sensitivity and bias in terms of reaction times and decision optimality metrics further substantiates the “disjoint mechanisms” model.

### Distinguishing mechanisms of endogenous attention from expectation

We have shown dissociable effects of endogenous spatial cueing on sensitivity and bias in our multialternative attention task. While enhanced sensitivity toward the cued location is a commonly reported effect of spatial attention (Bashinski & Bacharach, 1980; Ciaramitaro et al, 2001), do changes of choice criteria also reflect spatial attention’s effects (Summerfield & Egner, 2009)? Recent studies have suggested that attention includes processes that selectively alter decisional policies (e.g. criteria) based on priors and payoffs (Luo & Maunsell, 2018; Zénon & Krauzlis, 2012). Changes of choice criteria induced by spatial probabilistic cueing, therefore, likely reflect a key component of attention (Buschman & Kastner, 2015; Luo & Maunsell, 2018). Nevertheless, we evaluated the possibility that differences in event expectation, arising from different prior probabilities of events (cue validities) at the cued versus uncued locations, modulated choice bias independently of attention in our task.

First, we tested whether trivial strategies, based on event expectation (prior probabilities) alone, biased subjects’ choice outcomes when no signal evidence was available. We measured false alarm rates to each location on “no change” trials (Fig. 1E), which are indicative of each subject’s choice biases to the different locations in the absence of sensory evidence. We tested the distribution of these false alarm rates against three other distributions, each of which reflected alternative strategies by which choices could be influenced by priors alone: i) An optimal decision strategy to maximize the proportion of correct responses, based on signal expectation alone, which corresponds to consistently selecting the location with the greatest prior probability. Our data invalidated this hypothesis (p<0.001, goodness-of-fit test for multinomial data; indicating mismatch of model to data); ii) A probability matching strategy (Duda et al, 1973; Sugrue et al, 2005; Wozny et al, 2010)based on task specified priors, reflecting the relative proportion of change events at each location. In this case, the proportion of false alarms to each location should be distributed according to relative prior probabilities at that location (Figure Supplement 4C, unfilled bars with solid outline). Our data also invalidated this hypothesis (p=0.015; Figure Supplement 4C, compare ratio of heights of unfilled to filled bars at cued and uncued locations), and iii) A probability matching strategy based on perceived prior probabilities, reflecting the relative proportion of detections at each location for each subject on change trials. Perceived priors were quantified as the proportion of choices to the cued and uncued locations on all change trials, combined (sum of first 4 rows of the contingency table; Figure Supplement 4C, unfilled bars with dashed outlines). We tested whether the proportion of false alarms to each location were distributed according to these perceived priors. Our data invalidated this hypothesis also (p = 0.0012; randomization test). Overall, the mismatch of false alarm rates with any of these distributions indicates that expectation-based strategies did not influence choice bias in our task.

Next, we developed an alternative measure of spatial attention bias, unrelated to event expectation, to validate our m-ADC bias: the differential curvature of the risk, or utility, function (detailed description in Supplementary Methods, section *‘Risk curvature as a measure of bias’)*. Briefly, a higher curvature of the risk function at a location entails a greater penalty for sub-optimal placement of the criterion at that location, and vice versa (Fig. 5A). Therefore, subjects must place their choice criteria with greater precision (or care) at locations of higher risk curvature, to avoid significant penalties for perceptual decisions at those locations.

**Figure 5.**
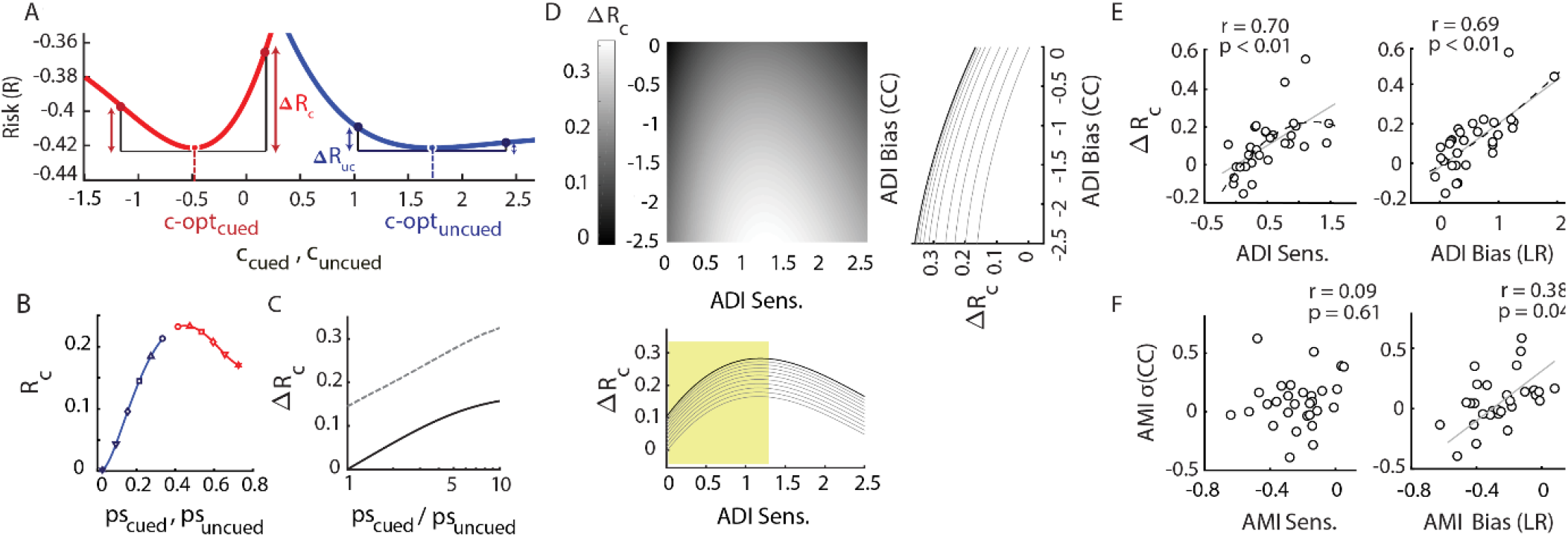
Covariation of sensitivity and bias with differential risk curvature. (**A-D**)Model, (**E-F**) Data. **A.** Risk as a function of criteria at the cued (red curve) and uncued (blue curve) locations (simulation). Red and blue dots: optimal criteria (c_opt_) at the cued and uncued locations, respectively. Colored arrows: increase in risk for a fixed deviation (increase or decrease) of the criterion from its optimal value at each location. **B.** Simulated variation of risk curvature (R_C_; second derivative of the risk function at the location of the optimal criterion) with signal probability at the cued (red) and uncued (blue) locations. Symbols on each curve: risk curvature values at each location, each pair corresponding to a different set of prior probabilities. **C.** Simulated variation of differential risk curvature (ΔR_C_; difference in risk curvature across the cued and uncued locations) with the ratio of signal probabilities at these locations. Solid line: d_cued_ = 1.00, d_uncued_ = 1.00; Dashed line: d_cued_ = 2.00, d_uncued_ = 1.00. **D.** Simulated variation of AR_C_with ADI bias (b_CC_) and ADI d’. Lighter shades: higher values of ΔR_C_. Insets: cross-section showing variation of ΔR_C_ with each parameter – ADI b_CC_, (right inset) and ADI d’ (lower inset) – at fixed values of the other parameter. **E.** Covariation of ΔR_C_, computed from behavioral data, with ADI d’ (left) and ADI bias b_LR_; right). Data points: individual subjects. Gray straight line: linear fit. Black dashed curve: quadratic fit. **F.** Covariation of AMI σ(CC) (modulation of the standard deviation of the choice criterion across cued and uncued locations) with AMI d’ (left) and AMI bias (bLR; right) respectively. Other conventions are as in panel E.

We simulated a multialternative attention task (Supplementary Methods) to examine the relationship between risk curvature, signal probability and bias. We found that risk curvature was higher at the cued location as compared to the uncued location (Fig. 5A) and varied systematically with the prior probabilities at each location (Fig. 5B). Differential risk curvature (ΔR_C_), the difference of risk curvature between the cued and uncued locations, varied monotonically with ratio of signal probabilities at cued and uncued locations (p_cued_/p_uncued_; Fig. 5C), despite identical sensitivities at these locations. These data indicate that differential risk curvature (ΔR_C_) is a measure of attentional bias: ΔR_C_ represents how much more precisely subjects must specify their choice criteria at the cued location versus at uncued locations. Finally, ΔR_C_ also varied monotonically with the difference of choice criteria across cued and uncued locations (ADI b_CC_; Fig. 5D, and right inset). On the other hand, differential risk curvature varied non-monotonically with the difference in d’ at cued and uncued locations (ADI d’; Fig. 5D, lower inset).

We tested whether a relationship between m-ADC choice criteria and the ΔR_C_ measure of spatial attention bias existed in our data. First, we measured ΔR_C_ across cued and uncued locations in individual participants (Supplementary Methods) and tested whether it was correlated with cueing-induced modulation of bias (both b_CC_ and b_LR_) and d’. We observed that ΔR_C_ was correlated with both bias and d’ modulations (ADI, Fig. 5E). The latter is an expected result because the range of Δd’ values in our data matched one (the rising) part of the non-monotonic curve shown in Figure 2-6 (shaded). Nevertheless, the data revealed evidence for a non-monotonic relationship at larger values of Δd’ (Fig. 5D, lower inset). Consequently, we tested whether any of these relationships would be better fit with a nonlinear function (second degree polynomial) using the normalized sum of squared residuals as well as adjusted R^2^ (coefficient of determination) values (Supplementary Methods). We observed that the normalized sum of squares residual measure was lower for a quadratic fit of ΔR_C_ vs. Δd’ (SS_err_: quadratic= 13.8×10^−2^; linear=14.5×10^−2^), whereas the measure was lower for a linear fit of ΔR_C_ vs. Δb (b_CC_: quadratic=16.0×10^−2^, linear=15.4×10^−2^; b_LR_: quadratic=11.2×10^−2^, linear=10.8×10^−2^). Confirming this trend, the adjusted R^2^ was greater for a quadratic (vs. a linear) fit for the variation with Δd’ (adj. R^2^: quadratic=0.37, linear=0.34), whereas the reverse was true for the variation with Δb (b_CC_: quadratic=0.27, linear=0.30; b_LR_: quadratic=0.49, linear=0.51). These results indicate a monotonic of variation of differential risk curvature with change in bias (both b_CC_ and b_LR_), but not with Δd’. We also performed multiple linear regression with ΔR_C_ as response variable and Δb and Δd’ as independent factors. We observed a higher standardized regression coefficient for Δb as compared to Δd’ (β: Δb=0.090, Δd’=0.066) as well a higher incremental R^2^ (R^2^_inc_: Δb=0.035, Δd’=0.018) further confirming a more robust, linear relationship of ΔR_C_ with m-ADC bias, rather than with d’.

Finally, we looked for yet another measure of attention bias that did not involve computing the risk curvature. We hypothesized that high attention bias would produce lower variation of the choice criterion, possibly due to the significant penalty associated with suboptimal positioning of the choice criterion at cued locations (Fig. 5A). Hence, we estimated, for individual participants, the standard deviation of the choice criterion across blocks at each location σ(CC), and tested for its correlations with modulations of sensitivity and m-ADC bias. We observed that σ(CC) modulations were significantly correlated with the modulations of bias (bLR; r=0.381, p= 0.038; Fig. 5F, right) but not with sensitivity (r= 0.097, p= 0.611; Fig. 5F, left).

These results demonstrate that m-ADC model estimates of bias, but not sensitivity, varied systematically and monotonically with two independent measures of spatial attention bias (ΔR_C_, σ(CC)) that were quantified from the data. Taken together, these lines of evidence strongly support the hypothesis that cueing induced changes of bias (m-ADC bias) in our attention task reflect spatial attention, rather than expectation, mechanisms. The alternative measures of attention bias (ΔR_C_ and σ(CC)) provide independent confirmation of our findings that sensitivity and bias are dissociable components of attention.

## Discussion

We demonstrate that perceptual (sensitivity) and decisional (bias) effects of endogenous cueing of attention operate through dissociable, rather than a single, unitary mechanism. Bias modulated systematically with endogenous cue validity (prior odds), was uncorrelated with sensitivity modulations, and correlated strongly with key decisional metrics including reaction times and decisional optimality indices. On the other hand, sensitivity enhancements were strongest at the cued location but not significantly different across uncued locations, suggesting that sensitivity control mechanisms allocate sensory processing resources in an all-or-none manner (“unitary spotlight”), whereas bias control mechanisms apportion decisional weights in a more graded manner.

Do sensitivity and bias changes in our attention task both reflect components of attention? There is emerging consensus that attention includes processes that selectively enhance sensory information processing (e.g. sensitivity) and also those that selectively alter decision policies for gating relevant information (e.g. choice criteria) based on sensory evidence, priors and/or payoffs (Buschman & Kastner, 2015; Dosher & Lu, 2000; Krauzlis et al, 2014; Eckstein et al, 2009; Luo & Maunsell, 2018). By this definition, changes of choice criteria induced by spatial probabilistic cueing do reflect a key component of attention (Buschman, 2015; Krauzlis et al, 2014; Luo & Maunsell, 2015). Our multialternative task, in conjunction with the m-ADC model, reveals that both bias changes, and sensitivity changes, independently contribute to performance benefits of endogenous attention.

Studies employing probabilistically-cued attention tasks without concomitant response probes (Cohen & Maunsell, 2009; Chanes et al, 2013; Luo & Maunsell, 2015) often seek to model the subject’s final decision in a multi-alternative task as arising from a conjunction of multiple one-dimensional decisions. This is not an appropriate formulation for the conventional attention task (Fig. 1B) because a) it assumes that the change/no change decision at each location is independent of the decisions at other locations; and, b) does not specify a decision strategy for trials in which decision variable values exceed criteria at multiple (more than one) locations. Other studies have sought to sidestep this problem by employing simpler models, but have had to factorize, *ad hoc*, or leave out some categories of responses in the contingency table (e.g. false alarms), based on assumptions that are not readily justified (Bashinski & Bacharach, 1980; Müller & Findlay, 1987).

Bachinski and Bacharach (1980) were among the earliest to test the effects of spatial cueing of attention on sensitivity and bias, using a dot stimulus detection task with probabilistic cueing (80% valid cues). They reported a benefit in sensitivity on the cued side, with little corresponding cost on the uncued side. Surprisingly, they found that bias was not different between cued and uncued locations. In direct contradiction to these findings, Shaw (Shaw, 1984) reported that, in luminance detection tasks, focusing attention produced bias (criterion) changes without concomitant changes of sensitivity. Along similar lines, Muller and Findlay (Müller & Findlay, 1987) demonstrated results similar to those of Shaw (Shaw, 1984), showing that attention primarily produced changes of bias at the cued (relative to uncued) location in luminance detection tasks.

These contradictions can be readily explained by shortcomings in the psychophysical models used to analyze these three-alternative task designs. Bashinski and Bacharach analyzed a three-alternative detection task with two one-dimensional models and incorrectly grouped misidentification (mislocalization) responses with misses (Figure Supplement 1). They also incorrectly partitioned the false-alarm rates according to an ad hoc rule, a pitfall highlighted by other studies as well (Luck et al, 1996; Müller & Findlay, 1987). Similarly, Muller and Findlay employed a two-stage signal detection model, which was unable to take into account all 9 stimulus-response contingencies in their three-alternative task., ignoring miss and misidentification responses though these constitute a significant proportion of overall responses for each event type (Fig.1E; Figure Supplement 1). In general, analyses that ignore any category of response can produce inaccurate estimates of sensitivities and biases (Figure Supplement 1).

To overcome these pitfalls, previous studies (Wyart et al, 2012, Luck et al, 1996, Downing, 1988) employed a post-hoc response probe paradigm in which subjects were cued to attend to one of multiple locations, and target presentation was followed by a response probe. Subjects had to indicate whether or not a target had appeared at the probe location, the cue being valid when the attentional cue matched the location of the response probe (response cue validity). Despite multiple potential stimulus locations, the response probe rendered this a 2-AFC (Yes/No) design that can be readily analyzed with independent one-dimensional signal detection models at each location. These studies have typically found systematic changes in sensitivity with response cue validity, but no reliable changes of bias (Luck et al, 1996; Wyart et al, 2012). In contrast, the m-ADC model revealed a graded variation of bias across cued and uncued locations, being highest at the cued location (Fig. 2C). Which result is correct?

These differences arise due to an essential distinction between response probe tasks and the detection and localization task, employed in this study. In the response probe task design, also employed in more recent neurophysiology studies (e.g. Luo & Maunsell, 2015; 2018) decisions need be based only on sensory evidence at the response probe location: there is no need to compare sensory evidence at the cued location against evidence at uncued locations. Therefore criteria in response probe tasks likely reflect signal expectation, rather than spatial attention mechanisms (Wyart et al, 2012, Summerfield & Egner, 2009; but see Rahnev et al, 2011). In contrast, in probabilistically cued attention tasks, employed in our study and in previous studies (e.g. Cohen & Maunsell, 2009) signal probability is the only relevant attention cue. Unlike response probe tasks in which independent Yes/No decisions are made at each location, our attention task requires subjects to detect and localize the stimulus (change) event by directly comparing sensory evidence across cued and uncued locations, on every trial. The modulation of m-ADC bias (difference in b_CC_ or b_LR_ across cued and uncued locations) by spatial cueing, therefore, reflects a decision policy that affords greater weight to signal information at the cued versus uncued locations, biasing the competition in favor of the sensory evidence at the cued location for making a localization decision on each trial. Consequently, choice criteria, as measured with the m-ADC model represent a measure of attention bias, consistent with the definition of attention that includes processes that selectively alter decision policies to benefit behavioral performance. The strong correlation between m-ADC choice criteria (b_CC_) with two different, data-driven measures of spatial attention bias (differential risk curvature and σ(CC)) further confirms that m-ADC choice criteria represent a measure of spatial attention bias.

Our findings and model are relevant, in general, for neuroscience studies that have reported a plethora of diverse, even conflicting, effects of attention on neural firing, tuning functions, neural correlations and neural synchrony; (Martinez-Trujillo & Treue, 2004; McAdams & Maunsell, 1999; Reynolds et al, 2000; Spitzer et al, 1988; Zénon & Krauzlis, 2012). A potential explanation for these diverse reports is that endogenous cueing of attention engages not a unitary process, but multiple processes, with different behavioral or neural signatures, which can be teased apart with the m-ADC model. For example, Chanes et al (2013) tested the roles of different frequencies of stimulation with rhythmic TMS over the right FEF, and reported frequency specific effects of TMS on sensitivity and criterion. They adopted a three-alternative task design in which subjects had to detect and report the location of a Gabor grating with one of three button presses (‘left’, ‘right’, ‘neither’). Again, because of the lack of an appropriate psychophysical model for analyzing this three alternative task, mislocalizations were entirely removed from the analysis as “error” responses, a pitfall that could lead to incorrect estimates of sensitivity and bias. Similarly, Luo and Maunsell (Luo & Maunsell, 2015) sought to determine if attentional modulation of neural activity in area V4 was correlated with sensitivity or bias changes while the monkey performed either a conventional three-alternative attention task (their Fig. 1A) or a response probe-type task. The authors concluded that neural activity signatures, including an increase in the firing rate of V4 neurons, were correlated with sensitivity (signal-to-noise ratio) changes, but not with criterion changes. As before, sensitivity and bias were quantified with a combination of one-dimensional models for the three-alternative task, or with a 2-AFC model for the response probe task, both of which, as indicated above, are not appropriate for measuring attention bias. Interestingly, Baruni et al (Baruni et al, 2015) reported results in direct contradiction to these findings. These authors manipulated absolute and relative rewards across different stimulus locations, and reported that neural modulation of V4 activity did not reflect the action of signal to noise (sensitivity) mechanisms. Again, these contradictions could potentially be reconciled if the m-ADC task and model were employed to analyze animals’ behavior in these attention tasks.

It is increasingly clear that cognitive processes, like attention or decision-making, are not unitary constructs. Psychophysical models, like the m-ADC model, are essential to tease apart component processes of these phenomena for understanding how cognitive processes emerge from neural activity and influence behavior, both in health and in disease.

## Materials and Methods

### Participants

37 subjects (21 males, age range 19-60 yrs.; median age 22 yrs.) with no known history of neurological disorders, and with normal or corrected-normal vision participated in the experiment. All participants provided written informed consent, and all experimental procedures were approved by the Institute Human Ethics Committee at the Indian Institute of Science, Bangalore. Seven subjects were excluded from analysis because of improper gaze fixation (see section on eye tracking, Supplementary Methods). Data from 30 subjects were included in the final analysis.

### Task

Participants were tested on a cued, five-alternative change detection task (Fig. 1A). Subjects were seated in an isolated room, with head positioned on a chin rest 60 cm from the center of a contrast calibrated visual display (22-inch LG LCD monitor, X-Rite i1 Spectrophotometer). Stimuli were programmed with Psychtoolbox (version 3.0.11;(Brainard, 1997)) using MATLAB R2014b (Natick, MA). Responses were recorded with an RB-840 response box (Cedrus Inc., California, USA). Subjects were instructed to maintain fixation on a central fixation cross during the experiment. Fixation was monitored with a 60 Hz infrared eyetracker (Gazepoint GP3). Subjects began the task by fixating on a fixation cross at the center of a grey screen (0.5° diameter). Following 200 ms, four full contrast Gabor patches appeared, one in each visual quadrant (Fig. 1A) at a distance of 3.5° from the fixation cross (Gabor s.d.= 0.6°; grating spatial frequency, 2 cycles/degree). The orientation of each Gabor patch was drawn from a uniform random distribution, independently of the other patches, and pseudorandomized across trials. After another 500 ms, a central cue (directed line segment, 0.37° in length) appeared. After a variable delay (600 ms-2200 ms, drawn from an exponential distribution), the stimuli briefly disappeared (100 ms), and reappeared. Following reappearance either one of the four stimuli had changed in orientation, or none had changed. The subject had to indicate the location of change, or indicate “no change”, by pressing one of five buttons on the response box (configuration in Figure Supplement 1D).

We term trials in which a change in orientation occurred in one of the four patches as “change” trials, and trials in which no change in orientation occurred as “catch” or “no change” trials. 25% of all trials were no change trials, and the remaining 75% were change trials. We term the location toward which the cue was directed as the “cued” (C) location, the location diagonally opposite to the cued location, as the “opposite” (O) location, and two other locations as “adjacent-ipsilateral” (A-I) or “adjacent-contralateral” (A-C) locations, depending on whether they were in the same visual hemifield or opposite visual hemifield to the cued location (Fig. 1 D-E). Changes occurred at the cued location on two-thirds of the change trials, at the opposite location on one-sixth of the change trials, and at each of the adjacent locations on one-twelfth of the change trials. Thus, the cue had a conditional validity of 67% on change trials, and an overall validity of 50%. The experiment was run in six blocks of 48 trials each (total, 288 trials per subject), with no feedback. In an orienting session prior to the experiment, subjects completed 96 trials (two blocks) with explicit feedback provided at the end of each trial about the location of the change and the correctness of their response. Data from these “training” blocks were not used for further analyses.

Ten subjects were tested on a version of the task that incorporated neutrally cued blocks. In this task, subjects were tested on a total of 8 experimental blocks (48 trials each; total 384 trials), with 4 blocks comprising predictively cued trials (as before) and the remaining 4 blocks comprised neutrally cued trials. On neutrally cued trials, the cue was made up of four directed line segments, each pointing toward one of stimuli in each of the four quadrants (Fig. 1A) and changes were equally likely at all four locations. Subjects were informed by on-screen instructions before the beginning of each block as to whether it was a predictive cueing or neutral cueing block, and the order of blocks were counterbalanced and pseudorandomized across subjects. All other training and testing protocols remained the same as before.

## Data analysis

### Contingency tables

Subjects’ responses in the task were used to construct 5×5 stimulus-response contingency tables, with change locations relative to the cue on the rows and response locations relative to the cue on the columns; no change events and responses were represented in the last row and last column respectively. Thus, each contingency table comprised five categories of responses: hits, misses, false-alarms, mislocalizations (or misidentifications) and correct rejections (Figure Supplement 1A). Since six values of orientation changes were tested at each location, the contingency table contained 100 independent observations (24 hits, 72 misidentifications and 4 false-alarms; the last category of responses do not depend on orientation change magnitude).

### Psychometric and psychophysical functions

To compute the psychometric function (percent correct as a function of orientation change angle), we calculated the proportion of trials in which the subject detected and localized the change accurately; this was computed separately for each location (relative to the cue). Percent correct values, across all angles of orientation change, were fitted with a three-parameter sigmoid function to generate the psychometric function (Fig. 2A, top). Psychometric functions in 24/30 subjects were estimated with a set of six change angles spanning 2° to 90° ([2,10,15,25,45,90]°), presented in an interleaved, pseudorandomized order across locations. As the psychometric function tended to saturate around 45°, in the remaining 6/30 subjects psychometric functions were estimated with a set of six change angles spanning 2° to 45°. The analyses were repeated by excluding this latter set of 6 subjects, who were tested on the more limited range of angles, and in all cases we obtained results similar to those reported in the main text. A single, combined psychometric curve was generated by pooling contingencies across all subjects and computing the above metrics for the pooled data (Fig. 2A, top). False alarm and correct rejection rates were calculated based, respectively, on subjects’ incorrect and correct responses during no-change trials.

### Model fitting and prediction analyses

To compute the psychophysical function (sensitivity as a function of orientation change angle), individual subjects’ response contingencies were fitted with the m-ADC model described previously. We estimated sensitivities and criteria with maximum likelihood estimation (MLE), using a procedure described previously (Sridharan et al, 2014). The sensitivity is expected to change depending on the magnitude of stimulus strength (change angle value); hence, different sensitivity values were estimated for each change angle tested. On the other hand, the criterion at each location was estimated as a single, uniform value across change angles. As change angle magnitudes at each location were distributed pseudorandomly it is reasonable to expect that the subject could not anticipate and, hence, alter her/his criterion for different values of change angles. Thus, the model estimated 28 parameters (6 d’ values for each of the 4 locations, and 4 criteria) from 100 independent observations in the contingency table.

We simplified m-ADC model estimation further using the following approach: As the probability of change was identical across the adjacent-ipsilateral and adjacent-contralateral locations (Fig. 1B), we compared the parameters (sensitivities, criteria) estimated for these two locations. Sensitivities and criteria were strongly correlated (opp. vs. adj: d’:ρ=0.58, p=0.002; c: ρ=0.74, p=0.0001; Figure Supplement 2), and not significantly different between these locations (difference of ipsi. vs. contra medians: Δd’=0.07, Δc=0.03, p>0.5, Wilcoxon signed rank test). Thus, responses from cue ipsilateral and contralateral sides were averaged, and treated as a single “adjacent” contingency in the contingency table. Therefore, the m-ADC model estimated 21 parameters (6 d’ values for each of the 3 locations – cued, opposite and adjacent – and their corresponding 3 criteria) from 63 independent observations in the contingency table. In all of the subsequent analyses we report psychophysical parameters calculated for these three locations relative to the cue, treating the two cue-adjacent locations (A-I, A-C) as a single “adjacent” location (A).Occasionally, parameters are reported as a single value for uncued locations; these were based on their average values across all three uncued locations.

For the behavior prediction analysis, (Fig. 1F-I)model fitting was done with only subsets of contingencies (hits and false-alarms, hits and misses or missed and false-alarms), as described in the Results. The predicted contingency table was compared with the observed contingency table with a goodness-of-fit (randomization) test (see Methods: *Statistical tests, correlations and goodness-of-fit*.).

### Psychophysical function fits

The psychophysical function was generated by fitting the sensitivity values across angles at each location with a three-parameter Naka-Rushton function allowing asymptotic sensitivity (d_max_) and orientation change value at half-max (Δθ_50_) to vary as free parameters, keeping the slope parameter (n) fixed at 2; a value determined from pilot fits to the data (Herrmann et al, 2010). As before, a single combined psychophysical curve was generated by pooling contingencies across all subjects and computing the above metrics for the pooled data (Fig. 2B, top). Values of d_max_ and Δθ_50_ reported in the main text correspond to pooled psychophysical fits with jackknife error bars (Fig. 2B, bottom left and right). Tests of significant differences in dmax and Δθ**50** values across locations were performed with a bootstrap approach: by comparing the true differences in values against a null distribution of differences calculated from contingency tables obtained by randomly shuffling, 1000 times, the labels across stimulus and response locations (cued, opposite, adjacent).

### Computing psychophysical parameters and their modulations

After fitting the m-ADC model to the contingency table to estimate sensitivity and criteria we computed, at each location, the mean sensitivity (d’_av_; average value across change angles), as well as two measures of bias: the choice criterion (b_CC_= c-d’_av_/2), and the likelihood ratio (b_LR_; Supplementary Methods: Model Derivation, equation 12). Choice criterion measure of bias (bias(CC)) was quantified as the deviation of the subject’s criterion (detection threshold) from the optimal criterion, which occurs at the midpoint of the means of the signal and noise distributions for equal prior probabilities of signal and noise. The lower the choice criterion (b_CC_) at a location, the lower the value for the decision variable at which the subject chooses to indicate signal over noise at that location, and, consequently, the higher the choice bias (for signal events) at that location. On the other hand, the likelihood ratio measure of bias (b_LR_) was calculated as the ratio of the conditional probability density of the signal to the noise at the value of the choice criterion. The higher the likelihood ratio bias at a location the higher the choice bias (for signal events) at that location. Quantifying likelihood ratio bias becomes relevant if there are significant differences in sensitivity across locations, as would be expected in attention tasks (e.g. cued versus uncued). For Gaussian decision variable distributions, both measures of bias are closely related (b_LR_ = exp(-d’ *b_CC_); Supplementary Methods: Model Derivation, equation 12).

In addition, we computed two measures to quantify attentional modulation of these parameters induced by endogenous cueing: a difference index and a modulation index. The attention difference index for each parameter was computed as the difference between the values of the respective parameter at the cued and uncued locations (e.g. d’_ADI_ = d’_cued_ – d’_uncued_) whereas the attention modulation index was computed as the ratio of the difference of the parameters at the cued and uncued locations to their sum (e.g. d’_AMI_ = (d’_cued_ – d’uncued)/ (d’_cued_ + d’_uncued_)).

### Statistical tests, correlations and goodness-of-fit

Unless otherwise stated, pairwise comparisons of different parameter values (e.g. mean d’ or bias) across locations was performed with the non-parametric Wilcoxon signed rank test with FDR (Benjamini Hochberg) correction for multiple comparisons. Correlations across different parameter values or modulation indices were typically Pearson correlations. For the optimal behavior, choice theory and reaction time analyses, robust correlations (“percentage-bend” correlations) were computed to prevent outlier data points dominating the correlations (Wilcox et al, 1994). Goodness-of-fit of the model to the data was assessed using a randomization test based on the chi-squared statistic; the procedure is described in detail elsewhere (Sridharan et al, 2017). A small p-value (e.g. p<0.05) for the goodness-of-fit statistic indicates that the observations deviated significantly from the model.

To test if there was covariation between sensitivity and bias within each subject’s data, we adopted the following procedure: Two contingency tables were constructed for each subject, with responses drawn from the first half of her/his respective session (first one half of the trials) and the second half (last one half of trials). Psychophysical parameters were estimated from these two subsets of data yielding two measures of each psychophysical parameter per subject. An n-way ANOVA analysis was performed with each measure of bias (b_CC_, b_LR_) as the response variable and sensitivity as a continuous predictor, with subjects as random effects.

### Analysis of reaction times

Reaction times were computed as the time from change onset to the time of response; no limit was placed on permitted response intervals, but subjects were asked to respond as quickly as possible. For each subject, trials in which RTs fell outside three standard deviations from the mean RT was considered outliers and excluded from further analysis. Each subjects’ mean RT at each location was normalized by dividing by the median RT at the cued location across the population, and correlated with psychophysical parameter estimates at the respective location. Partial correlations were performed to identify dependencies between RT and measures of bias while controlling for the effects of d’, and vice versa. We also performed multilinear regression analysis with RT as the response variable and change angle, mean sensitivity and bias at each location as predictors; all predictors were scaled to zero mean and unit variance before the analysis. Regression coefficient magnitudes for sensitivity (βd’) and bias (βb-LR) were compared with a bootstrap test; as our hypothesis was that the regression coefficient for bias would be higher than that for d’ because of the similar graded variation of bias and RT across locations, we constructed a null distribution of regression coefficient differences by shuffling the location labels for the d’ and bias values randomly and independently across subjects. In order to discount the hypothesis of motor bias, robust correlations were computed between mean RTs and mean FA rates for each experimental block after subtracting the mean RT and FA values subject-wise to account for subject specific effects. An n-way ANOVA was applied, as before, treating RT as the response variable with FA rates as continuous predictors, and subjects as random factors.

Further details regarding eye-tracking, model comparison analysis, and analyses of optimal decisions and risk curvature are presented in Supplementary Methods.

## Acknowledgments

The authors would like to thank Kelsey Clark, Behrad Noudoost and Nicholas Steinmetzfor their comments on a preliminary version of this manuscript, and Subbulakshmi Sankaranarayanan for help with data collection. This research was funded by a Wellcome Trust-Department of Biotechnology India Alliance Intermediate fellowship, a Science and Engineering Research Board Early Career award, a Pratiksha Trust Young Investigator award, a Department of Biotechnology-Indian Institute of Science Partnership Program grant, and a Tata Trusts grant (all to DS).

## Supplementary Information

### Supplementary Methods

#### Eye-tracking

Subjects’ gaze was binocularly tracked and the deviation in their gaze from the fixation cross was recorded and stored in degrees. Trials in which the eye-position deviated by more than 2 degrees from fixation either in the x- or in the y-direction, from the onset of the Gabors until the final response, were removed from further analysis. All of our subjects were South Asian and several exhibited dark pigmentation of the iris, rendering it nearly indistinguishable from the pupil. Hence, the contrast of the pupil (relative to the iris) was weak, and the tracker occasionally lost the location of the pupil; trials in which this occurred for more than 100 ms continuously were also excluded from the analysis. Finally, we excluded all subjects for whom the combined rejection rate (from eye deviation and lost tracking) was more than 30% (7/37 subjects). Among the set of subjects who remained (n=30), the rejection rate was 8.6% [4.2-17.7%]. We also confirmed with a randomization test, based on the chi-squared statistic (see below), that these rejected trials did not significantly alter the distribution of responses in the contingency table for any subject (p-values for distributions before and after rejection: 0.99, median across subjects).

#### Model comparison analysis

##### i. Comparison with models assuming identical parameter values at uncued locations

We compared the goodness of fit of our default model with two other models, one that assumed equal sensitivities at all uncued locations (m-ADC_eq-d_ model; d’_opp_ = d’_adj_) and one that assumed equal criteria at all uncued locations (m-ADC_eq-c_ model; c_opp_ = c_adj_). Models were compared with the Akaike Information Criterion (AIC), that represents a tradeoff between model complexity (the number of fitted model parameters) and goodness-of-fit, (based on the log-likelihood function); a lower AIC score represents a better candidate model. Similar results were obtained when using a Bayesian Information Criterion (BIC). Because d’ was estimated separately for each of the six change angles, the number of fitted parameters in the m-ADC_eq-d_ model reduced to 15 (from 21 for the standard m-ADC model) and to 20 in the m-ADC_eq-c_ model.

##### ii Comparison with 2AFC model combination across locations

We compared the performance of the 4-ADC model with that of a combination of four one-dimensional SDT models (2-AFC or Yes/No models).2-AFC parameter estimates for each subject were obtained by separating their responses into four 2×2 contingency tables, one for each location, including only hits, misses, false-alarm and correct rejections; mis-localization responses were excluded because these cannot be categorized into any particular category of 2-AFC response. Each contingency table was then separately fit with a one-dimensional SDT model. Data from each subject’s experimental session was split into two sets, comprising the first and second half of the data acquired from that session. First, d’ and bias parameters were estimated for each of the split-halves and parameters estimated from the first half were correlated with those from the second half using robust correlations. Second, we performed a prediction analysis based on the same split-half analysis of the data. Briefly, for each subject, we estimated model parameters using each model from each half of the data and estimated response probabilities in the contingency table based on these model parameters. Next, we correlated modeled response probabilities from the first half against the observed response probabilities in the second half, and vice versa. These correlations represent a measure of cross-validation (or prediction) accuracy of each model. Because 2-AFC model parameters directly fit the data without additional degrees of freedom (Sridharan et al, 2014; 2017), 2-AFC model correlation values are equivalent to those obtained by directly correlating observed response probabilities across the split-halves.

#### Analysis of optimal decisions

Optimal decisions, in the m-ADC model framework, seek to minimize risk (Supplementary Methods: Model Derivation, equation 3). Under certain assumptions (e.g. C^l^_k_ = C^l^_m_, m ≠ l or for a given stimulus event type, the cost of all types of response error are identical)the optimal decision surfaces comprise hyperplanes in the multidimensional decision space. Our model fitting analysis suggests that observers choice criteria were closely in line with this optimal family of decision surfaces (as defined in Supplementary Methods: Model Derivation, equation 12, 13 and 14).

One potential definition of optimality is when the subject seeks to maximize the number of correct responses and minimize the number of errors. In this case, cost ratio at location j, β_obs-j_ = (C^0^_0_ − C^0^_j_) / (C^j^_j_ − C^j^_0_) = (C_CR_ − C_FA_)/ (C_Hit_ − C_Miss_) = 1 (where C^x^_y_ is the cost of responding to location x when the event occurred at location y). Yet, we noticed that β_j_, calculated as β_j_= (p_j_/p_i_)/b_LR-j_ (Supplementary Methods: Model Derivation, equation 8, 10 and 11), systematically deviated from 1 across the population of subjects (Fig. 4F), suggesting that subjects did not assume an equal relative cost for the two kinds of error responses (false alarm vs. correct reject and misses vs. hits). We also noticed that β_obs_ was different for the different locations, being predominantly greater than 1 at the cued location and less than 1 at uncued locations (Fig. 4F). Since there is only one type of correct rejection response, and since the model assumes that the cost for false alarms to each location is identical (Supplementary Methods: Model Derivation, equation 7), subjects assuming a different relative cost of hits to misses at the different locations (cued, opposite, adjacent) would be a somewhat implausible assumption. Rather, we propose that the reason for the difference in β_obs_ across the different locations is because the subject assumed a single, common cost ratio (β^s^_opt_) at all locations, but deviated from this (subjectively) optimal ratio at some locations where she/he did not consider it necessary to perform optimally. To determine this subject-specific β^s^_opt_ we tested different values of β and selected the one that minimized the deviation of the actual risk from the optimal Bayes risk assuming a single β^s^_opt_ at all locations (ΔRisk^obs-opt^); again this deviation is zero if the subject did not deviate from β_opt_ at any location (ΔRisk^obs-opt^ = Σ_i_Σ_j_C^i^_j_p_obs_^j^_i_ – p_opt_^j^_i_). Following some algebra, it is clear that C^i^_j_ for hits and misses (or misidentifications) is a function of β_opt_, C_FA_ and C_CR_ (see Supplementary Methods: Model Derivation, equations 6, 8 and 9, along with the following assumptions: a) C^j^_j_ = C^0^_0_β_opt_; b) C^j^_0_ = C^0^_j_/β_opt_; also C^j^_i_ = C^j^_0_ for all i ≠ j) whereas p_obs_^j^_i_ is a function of β_j_ and d_i_, and p_opt_^j^_i_ is a function of β_opt_ and d_i_.

For each subject three indices of optimal performance were computed. Two indices quantified the optimality of overall performance: i) an objective sub-optimality index (SI_O_) defined as the deviation of the subject’s cost ratio from an objectively optimal cost ratio (β_opt_=1) for maximizing correct responses, and computed as the magnitude of the logarithm of the ratio of β^s^_opt_ to β_opt_ (SI_O_= |log(β^S^_opt_ /β_opt_)|)and ii) a global sub-optimality index (SI_G_), defined and computed as the magnitude of the deviation of the optimal Bayes risk from the actual risk (SI_G_ = ΔRisk^obs-opt^). A third, locational sub-optimality index was defined as the deviation of the cost ratio at each location from the subject’s own optimal cost ratio, and computed for each location(SI_L_) as the magnitude of the logarithm of ratio of β_obs_ (observed cost ratio) at that location and the subject’s own β^s^_opt_ (SI_L_ = |log (β_obs_/β^s^_opt_)|). These sub-optimality indices were correlated with sensitivities, biases and their modulations using robust correlations.

#### Risk curvature as a measure of bias

We posit that subjects, when they have some form of prior information about events at different locations, would afford greater attentional bias to a location if greater penalties for incorrect (or sub-optimal) decisions occurred there as compared to other locations. Therefore, we propose that the differential curvature of the risk (or utility) function (ΔR_C_)is a measure of this attentional bias. The risk measures the total average cost associated with a specific decision strategy, given sensory evidence, priors and payoffs (Supplementary Methods: Model Derivation, equation 11-12). The curvature of this risk (R_C_) measures the “sharpness” with which the risk curve rises on either side of the optimum criterion. Briefly, a higher curvature of the risk function at a location entails a greater penalty (increase in risk) for sub-optimal placement of the choice criterion at that location, where the subjects must place their criterion with comparatively greater precision (e.g. Fig. 5A). We compute the curvature (second derivative) rather than the slope (first derivative) for measuring this bias, because the slope of the risk curve at the optimal criterion is zero at all locations. In the probabilistically cued attention task, the cued location is more important for decisions than uncued locations. Consequently, a higher value of risk curvature should occur at the cued location (Fig. 5B, red; simulated data, see below) as compared to other, uncued (irrelevant) locations (Fig.5B, blue) and this difference between the magnitude of the risk curvatures (Differential risk curvature or ΔR_C_) can be interpreted as a measure of attentional bias towards the cued location.

Therefore, we propose that processes linked to attention may control the precision with which choice criteria must be specified at cued versus uncued locations, considering the differences in risk profiles at each location. We expect the ΔR_C_ bias measure to be unrelated to signal expectation because while expectation is related to stimulus statistics (e.g. signal probability), we consider it unlikely to be associated with the evaluation of differential risks or penalties at the cued and uncued locations.

#### Risk curvature analysis for simulated attention task (Fig. 5A-D)

We simulated an m-ADC task with two potential stimulus locations. We denote the signal probability at cued and uncued locations as p_s-cued_ and p_s-uncued_, and the probability of no change as p*φ*. The cost ratio, β, was set to unity for both locations, corresponding to a goal of maximizing percent correct. To rule out the effects of sensitivity in these simulations, d’ at cued and uncued locations were set to identical values (d’_cued_ = d’_uncued_ = 1.0). In order to calculate risk curvature, we first determined the optimal criterion for both cued and uncued location using the relationship between criteria, sensitivities, priors and payoffs for the m-ADC model (Supplementary Methods: Model Derivation, equation 11). Using these sensitivity and criterion values, we generated the conditional response probabilities using m-ADC model equations. The risk (or utility) at for each stimulus response contingency was calculated as the product of the cost and probability of that contingency. The total risk (R) was then determined as the sum of the risks for all stimulus response-contingencies (Σ_i_Σ_j_C^i^_j_p_obs_^j^_i_) and across all locations. To determine risk curvature (R_C_), we varied the choice criterion about its optimum at each location across a range of values (±1.0), calculated the risk function at each value of the choice criterion, and then computed it’s second derivative. ΔR_C_ was calculated as the R_C_ difference across cued and uncued locations.

#### Relationship between m-ADC choice criteria and alternative measures of spatial attention bias

To test for correlation between m-ADC decision criteria and differential risk curvature (ΔR_c_) in our data, first we estimated sensitivity, criteria and ΔR_C_ for individual participants, using the same procedure as described above. Because our task lacked an explicit payoff structure, for these analyses we calculated the risk based on the observer-specific subjective cost ratio (β^s^_opt_), as reported in the decision optimality analyses (assuming a uniform cost ratio (β=1.0) produced similar results).

We tested whether ΔR_C_ was correlated with cue-induced modulation of bias (both CC and LR) and d’, using robust correlations (percentage-bend). To see if these relationships would be better fit with a quadratic function (second degree polynomial) relative to the linear fit, we determined normalized sum of squared residuals for each fit, calculated as sse/(n-m-1), where sse is the sum of squared residual of fit; n is the number of samples or data points and m is the order of the polynomial (lower value represents better fit of the model to data). Quadratic and linear fits were also compared using adjusted R^2^ (coefficient of determination) values, which measures how well the predictors explain the data while controlling for the number of predictors. A higher adjusted R^2^ is indicative of more informative predictors. We further performed a multiple linear regression analysis, with ΔR_C_ as the response variable and ADI-bias (b_LR_) and ADI-d’ as independent factors. In order to determine which of these factors exhibited a stronger relationship with ΔR_C_, we computed incremental R^2^ values associated with each factor, by calculating the increase in adjusted R^2^ upon adding each factor into a multiple regression model. A larger value of incremental R^2^ for a factor indicates a stronger linear relationship with the response variable.

Finally, we computed another measure of attentional bias that did not involve computing the risk curvature. For each participant and each location, we calculated σ(CC): the standard deviation of the choice criterion (b_CC_) values estimated with a jackknife approach. We correlated the AMI index of σ(CC) against the AMI indices of d’ and bias (b_LR_). All correlations were performed using robust correlations.

## Supplementary Figures

**Figure Supplement 1.**
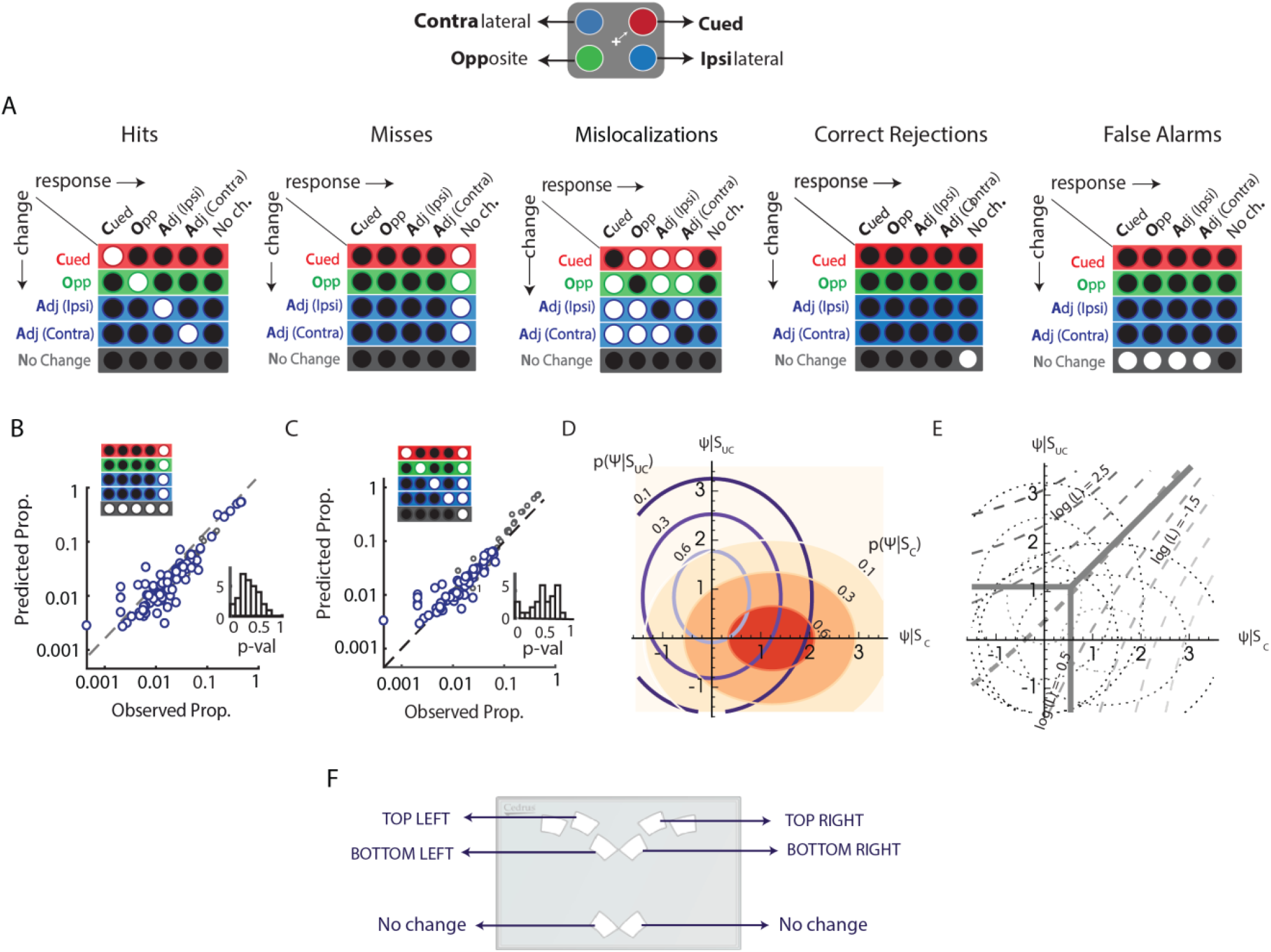
Stimulus-response contingencies, m-ADC model schematic and response schematic for the probabilistically cued attention task (Fig. 1A, main text). **A.** Schematics showing the different stimulus response contingencies: left to right - ‘Hits’ – response indicating the correct location of change; ‘misses’: no-change response on change trials; ‘mislocalization’: response indicating one location when the change happened at another location; ‘correct rejections’: correct no-change response, and ‘false alarms’: response indicating some location of change during no-change trials. **B.** Same as in Fig. 1G (main text), but showing predictions based on misses and false-alarms alone. Other conventions are as in Fig. 1G (main text). **C.** Same as in Fig. 1G (main text), but showing predictions based on hits and misses alone. Other conventions are as in Fig. 1G (main text). **D.** Contours of bivariate decision variable distributions (joint probability density function, pdf) in a task employing the method of constant stimuli in which stimuli or events (e.g. changes) can occur at one of multiple different strengths at the cued location (x-axis) or at an uncued location (y-axis). In the task shown in Fig. 1A (main text) this corresponds to different magnitudes of orientation changes, that are employed to measure psychometric functions as shown in Fig. 2A (main text). Red, filled contours: pdf contours for changes at the cued location; blue, unfilled contours: pdf contours for changes at an uncued location. Plots are based on average psychophysical functions calculated from experimental data (Fig. 2B; main text). **E.** Optimal decision surfaces in the m-ADC model. Dashedlines: Contours of constant log-likelihood ratio (log (L)) of the two decision variable distributions shown in panel D. Thick solid lines: Planar decision surfaces in the m-ADC model. The oblique decision surface closely approximates the optimal decision contours of constant log-likelihood. Thick dashed line: Representative optimal decision contour (log(L) = 0.5). Dotted lines: Decision variable pdf contours from panel D, shown in the background, for reference. Plots are based on average criteria calculated from experimental data (Fig. 2B; main text). **F.** Schematic showing the arrangement of buttons in the response pad (CedrusTM – RB 830). The four top inner buttons correspond to the four possible locations of change. Either of the bottom two buttons can be pressed to indicate no change.

**Figure Supplement 2.**
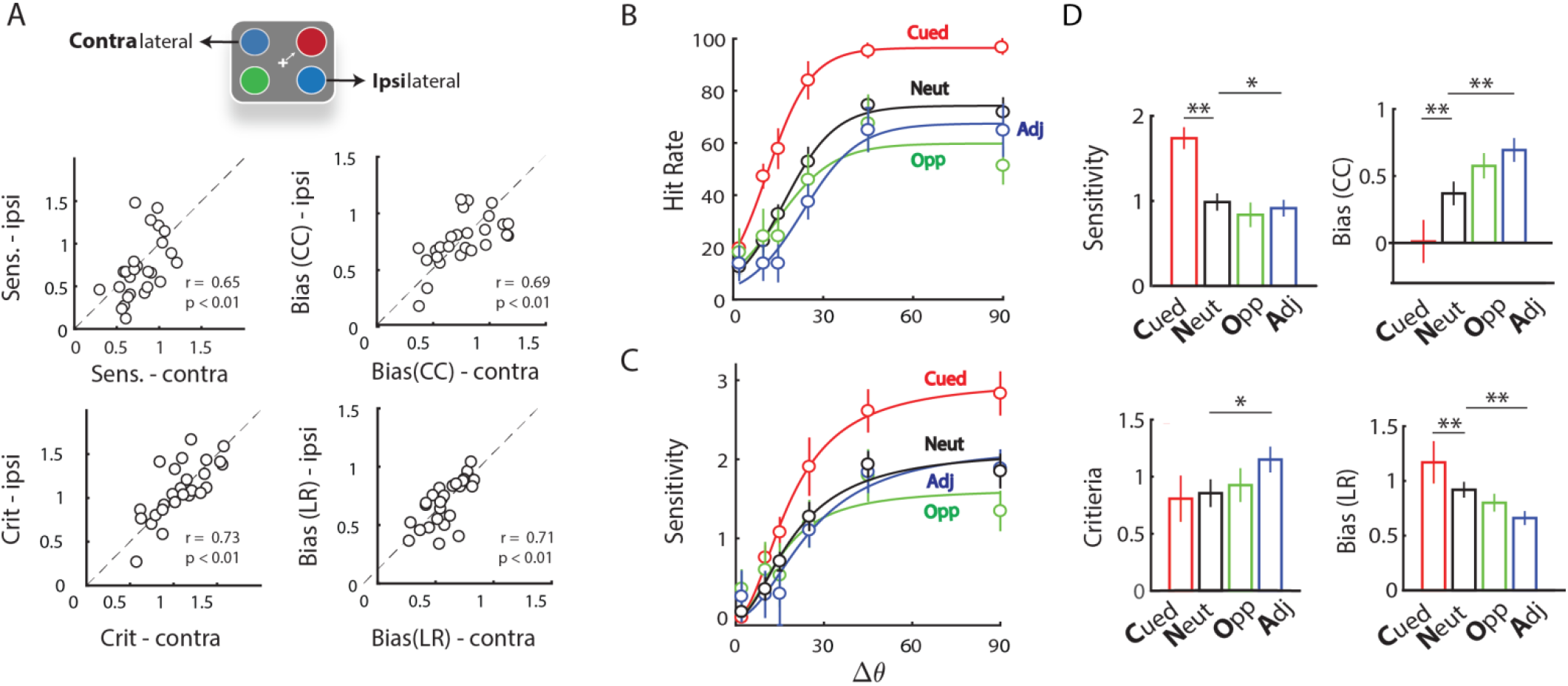
Comparison of parameters at cue-ipsilateral and cue-contralateral locations and sensitivity and bias effects in neutrally-cued and predictively-cued blocks. **A.** Correlations between sensitivity (top left), criteria (bottom left), bias (b_CC_) (top right) and bias (b_LR_) (bottom right) values for ipsilateral and contralateral sides. Data points represent subjects (n=30), and diagonal shows line of equality. **B.** Average psychometric function (n=10 subjects, who performed both the neutrally cued and predictively cued blocks) showing hit rates at different change angles (Δθ) at the cued, opposite and adjacent locations and for neutrally cued trials across all locations (black). Circles, curves and error bars follow the same convention as Fig. 2A. **C.** Same as in panel B, but psychophysical functions showing sensitivity at different change angles. Circles, curves and error bars follow the same convention as Fig. 2B. **D.** Median sensitivity, criterion, choice criterion bias (b_CC_) and likelihood ratio bias (b_LR_) measures of bias at cued, opposite and adjacent locations and for neutrally cued trials across locations. Other conventions are same as Fig. 2C (main text).

**Figure Supplement 3.**
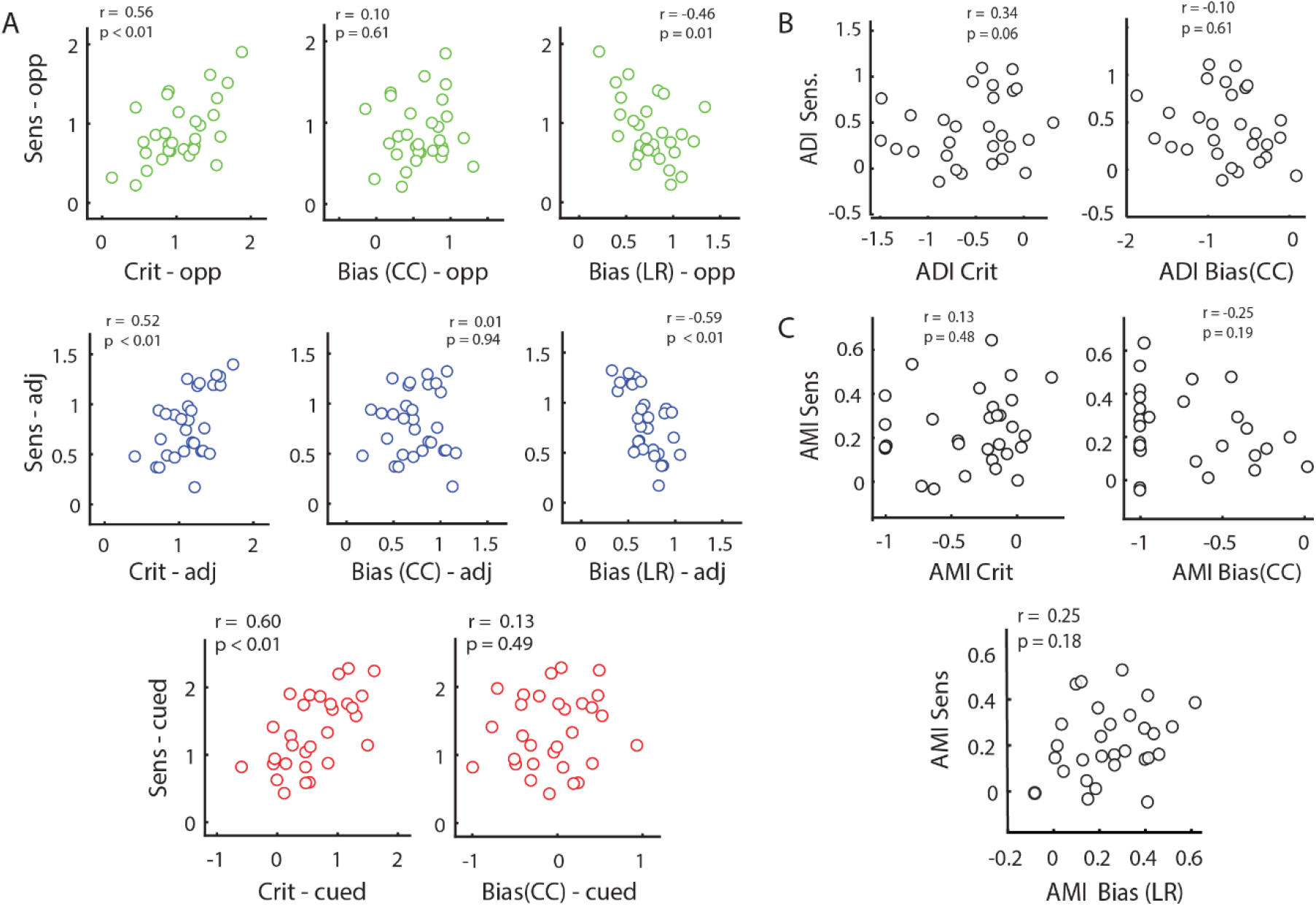
Covariation among psychophysical parameters. **A.** (Top row) Covariation of sensitivity with criterion, choice criterion bias (bCC) and likelihood ratio bias (bLR) at the cue-opposite location. (Middle row) Same as top row, but for cue-adjacent location. (Bottom row) Covariation of sensitivity with criterion and bCC at cued location. Color conventions are as in Fig 1E (main text). **B.** Covariation of difference indices (ADI) of sensitivity with those of criteria (left) and bCC (right). **C.** Covariation of modulation indices (AMI) of sensitivity with that of criteria (upper left), bCC (upper right) and bLR (lower).

**Figure Supplement 4.**
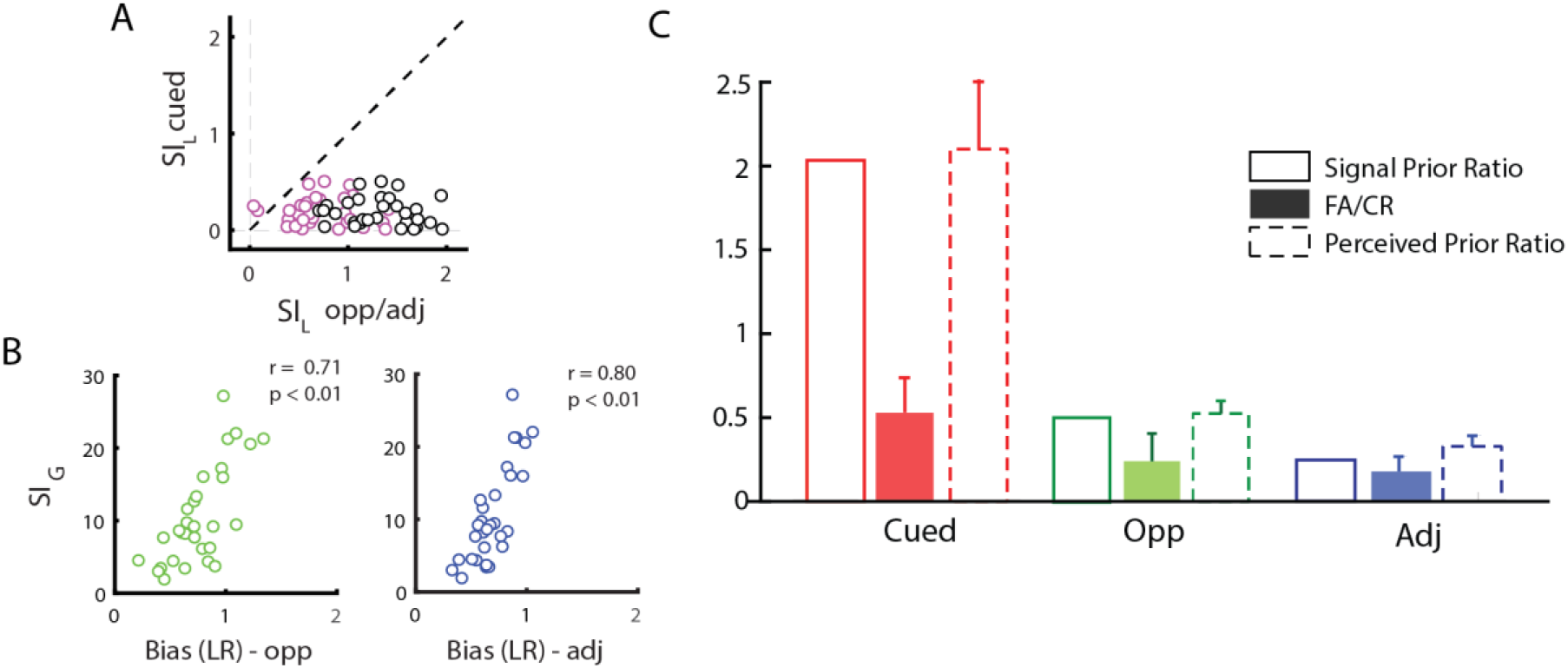
Covariation of sensitivity and bias with optimal decision indices and effect of priors on choice outcomes. **A.** Covariation of the locational sub-optimality index (SI_L_) at the cued (y-axis) and uncued (x-axis) locations. Black: SI_L_ at cued versus opposite locations; magenta: SI_L_ at cued versus adjacent locations. Dashed diagonal line: line of equality. Data points: Individual subjects. **B.** Covariation of the global sub-optimality index (SI_G_) with bias at the opposite (left) or adjacent (right) locations. Data points: Individual subjects. **C.** Effects of event prior probabilities on choice outcomes during no change trials. Filled bars: The proportion of false alarms at the cued, opposite and adjacent locations. Unfilled bars, solid outlines: Task-imposed signal priors. Unfilled bars, dashed outlines: Perceived priors. These values are plotted as a ratio over the proportion of correct rejections (FA:CR ratio), no-change trials (signal prior ratios) or no-change responses (perceived prior ratios). Error bars: s.e.m. Color conventions are as in Fig. 2A (main text).

## Supplementary Methods: Model Derivation

### mADC decision rule for a multialternative attention task employing the ‘method of constant stimuli’

The conventional attention task shown in Fig. 1A involves ‘M+1’ stimulus events: changes at one of ‘M’ locations or no change.

We denote the change event at each location as *S*_1_,…, *S_M_*, and the no change event as *S*_0_. Similarly, we denote the event (or response) locations with indices 1,2,…, *M* and denote the no change event (or response) with the index 0 (or *ϕ*). Finally, we denote *p_k_* as the prior probability of change at location *k* and *p*_0_ as the prior probability of no change.

Extending one dimensional signal detection models, we define a multivariate decision variable, **Ψ**, whose components Ψ_*j*_ denote the scalar decision variable at each location *j* where *j* ∈ {1,2,…, *M*}.

We derive the optimal decision rule for the case where the change at each location *k* can occur at one of *N_k_* different strengths (change magnitudes). The derivation is based on Bayesian decision theory for the optimal detection of signals in noise (Middleton and van Meter, 1955).

The observer makes a decision/response *γ_k_* to one of *M* locations *k* or no response 0. This is denoted by the variable *δ*(*γ_k_*|**Ψ**), such that

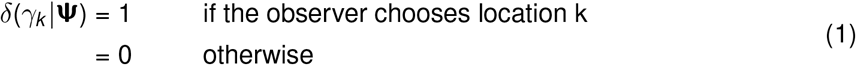

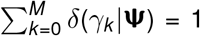, since the observer can choose only one of the *m* locations (or 0 in case of no change).

In this case, the average Bayesian risk is given by:

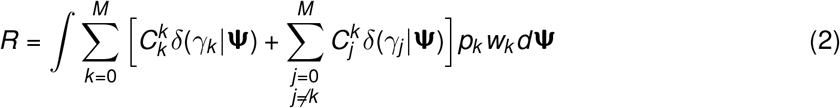

where 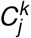 is the cost of indicating a response to location *j* when the change occurred at location *k*, where *k*, *j* = 0, 1, 2,…, *M* and *w_k_* = *p*(**Ψ**|*S_k_*), i.e. the conditional probability density of the decision variable given that a change occurred at location *k*. Grouping terms by *δ*(*γ_k_*|**Ψ**),

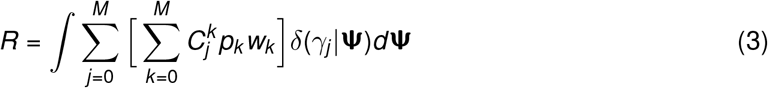

Since only one of the *δ*(*γ_j_* |**Ψ**) = 0, our objective is to find the *j* that minimizes the sum 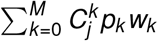. Let us consider the *N_k_* magnitudes of changes associated with events *S_k_* at each location *k*, and posit that each occurs with prior probability 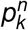 and with conditional density 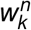. Thus,

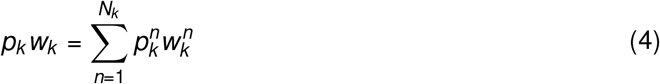

We assume also that costs are not different for the different magnitudes of change at each location. The sum to be minimized can then be written as:

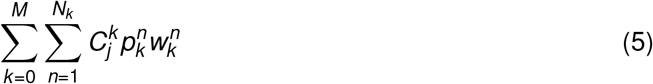

Let us consider choosing between two alternatives *i* & *j*. The choice between *i* & *j* depends on the relative average risks, given by:

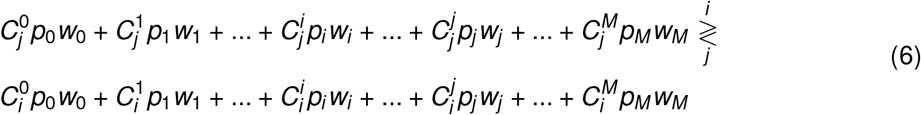

where the notation 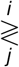 means choose *i* if LHS>RHS, and choose *j* otherwise.

We assume

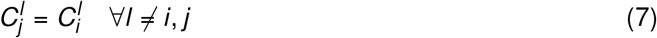

implying that the cost of making an incorrect response to any location (*i* or *j*) is the same for changes at a given location (*l*). Generally, we assume that for every event type *S_k_*, the cost of making an incorrect response is the same, although this cost may be different across different event types. With this simplification, equation (6) reduces to:

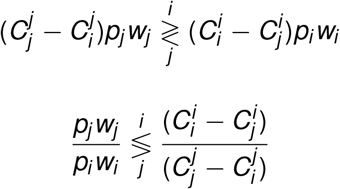

where, the reversal of inequality happened because 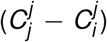 is defined to be non-zero and negative because the cost for making correct response is less than cost for making an incorrect response.

The left hand side of the previous equation is the generalized likelihood ratio, or the posterior odds ratio (product of the prior odds and the liklihood ratio). Introducing new notation for the cost ratio term on the right hand side,

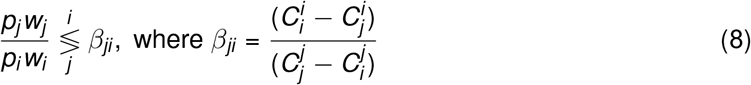

Substituting (4), this equation becomes

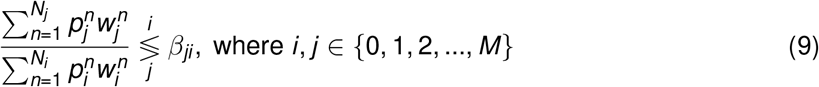

In our experiment, *N_i_* = *N_j_* = *N* (6 change angles in our task, Fig. 1A) and 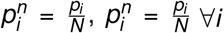, *j* ∈ 0, 1, 2,…, *M* (prior probabilities are evenly divided among the change angles). The only exception to this formula is for the no-change event for which *N*_0_ = 1 (no change events do not occur at different change magnitudes).

We first examine the case for making a decision regarding change at location *j* vs no change.

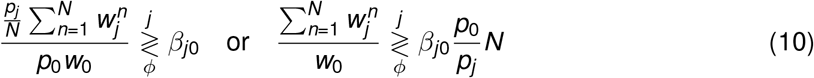

where *ϕ* denotes a no change response. We assume that 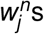 are multivariate Gaussians with mean 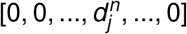 (an *m* × 1 vector and identity covariance matrix. Again, as a special case 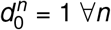; no change distribution has zero mean. Thus,

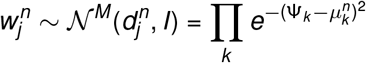

Now, equation (10) above simplifies to

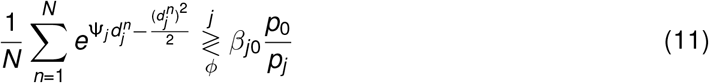

Therefore, optimal decision surfaces are hyperplanes of constant ψ_*j*_ even if, in this case there is no evident closed form solution for Ψ_*j*_. These hyperplanes correspond to the criteria, Ψ_*j*_ = *c_j_*. Consequently, the generalized likelihood ratio measure of bias for choices to location *j* is defined as:

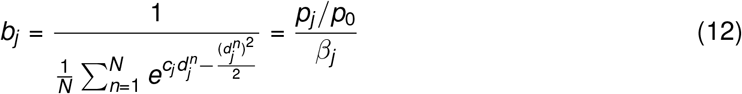

which is the harmonic mean of the biases calculated from individual sensitivity values for each change angle 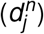.

We next examine the case for making a decision at location *j* vs location *i*.

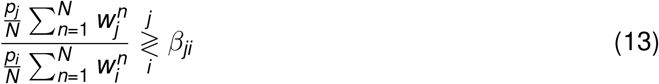

Dividing the numerator and denominator by *p*_0_ *w*_0_ & after some algebra,

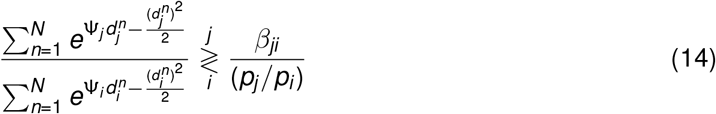

Thus, the optimal decision surface for choosing change at location *i* versus change at location *j* are not hyperplanes. However, planar decision surfaces provide excellent approximations to these optimal surfaces, as shown in Supplementary Fig. S2B, and as derived next.

### Demonstration that optimal decision surfaces for the choice between locations *i* & *j* can be closely approximated by planar surfaces

We apply the logarithm on both sides of equation (14):

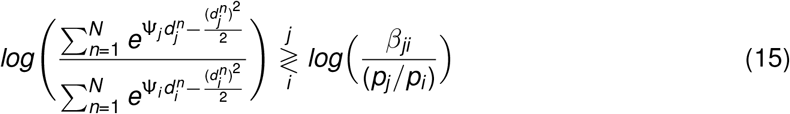

Since the numerator and denominator are identical in form, except for the location indices, we consider the numerator term alone 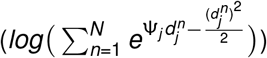, without loss of generality. We consider two cases, a) for small Ψ_*j*_ and b) for large Ψ_*j*_.

a) First, consider small 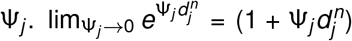, where, we have ignored terms 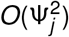 and higher in the expansion for 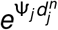. Thus, the numerator simplifies to:

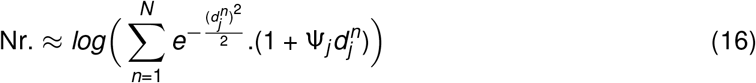

Further simplifying,

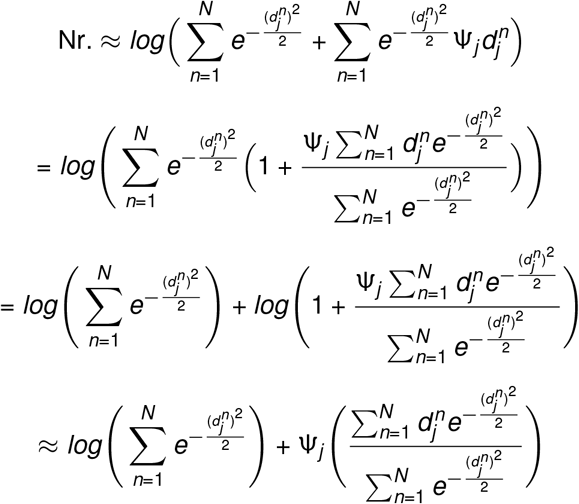

where, we have used the approximation *log*(1 + *x*) ≈ *x* for small *x*. This has the form,

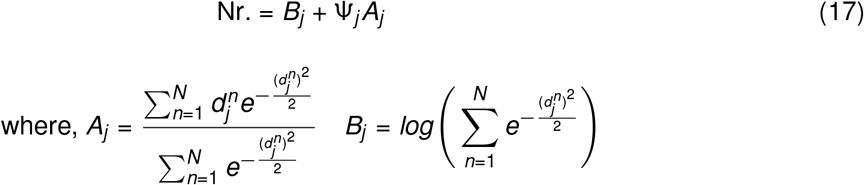

An identical form holds for the denominator term in equation (15) for small Ψ_*i*_.

b) Next, consider the numerator term from equation (15) for large Ψ_*j*_

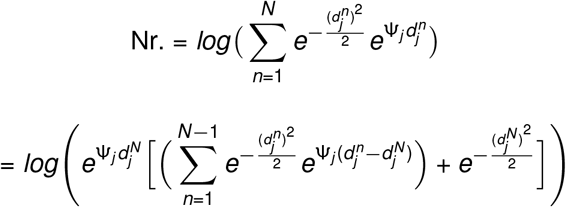

We consider 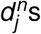 to be sorted such that 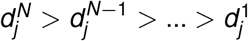, which represent to the sensitivities corresponding to different change magnitudes (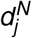, corresponding to the largest, and 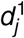 to the smallest change magnitude).

As 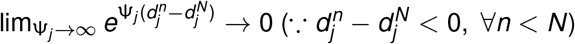, this simplifies to

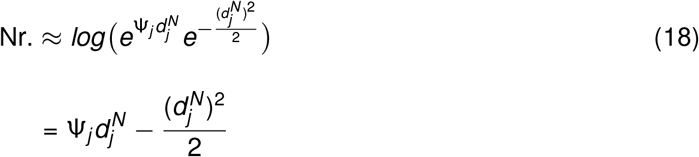

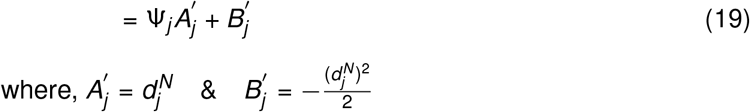

Again, an identical form holds for the denominator term in equation (15) for large Ψ_*i*_.

Finally, we compute the decision surfaces for the choice between locations *i* and *j* for the different combinations of Ψ_*i*_ and Ψ_*j*_ magnitudes.

For small Ψ_*i*_, large Ψ_*j*_, the decision surface are given by (equations (17) and (19)):

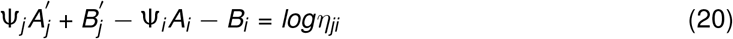

Similarly, for large Ψ_*j*_, large Ψ_*i*_, the decision surfaces are given by (equation (19)):

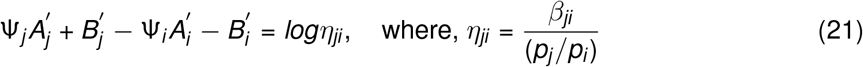

Decisions surfaces for the other two combinations (small Ψ_*j*_, large Ψ_*i*_ and small Ψ_*j*_, small Ψ_*i*_) may be similarly derived. In each case, the decision rule is approximated by a planar decision surface (decision hyperplane).

